# Genetic, transcriptomic, histological, and biochemical analysis of progressive supranuclear palsy implicates glial activation and novel risk genes

**DOI:** 10.1101/2023.11.09.565552

**Authors:** Kurt Farrell, Jack Humphrey, Timothy Chang, Yi Zhao, Yuk Yee Leung, Pavel P Kuksa, Vishakha Patil, Wan-Ping Lee, Amanda B. Kuzma, Otto Valladares, Laura B. Cantwell, Hui Wang, Ashvin Ravi, Claudia De Sanctis, Natalia Han, Thomas D. Christie, Kristen Whitney, Margaret M. Krassner, Hadley Walsh, SoongHo Kim, Diana Dangoor, Megan A. Iida, Alicia Casella, Ruth H. Walker, Melissa J. Nirenberg, Alan E. Renton, Bergan Babrowicz, Giovanni Coppola, Towfique Raj, Günter U. Höglinger, Lawrence I. Golbe, Huw R. Morris, John Hardy, Tamas Revesz, Tom T. Warner, Zane Jaunmuktane, Kin Y. Mok, Rosa Rademakers, Dennis W. Dickson, Owen A. Ross, Li-San Wang, Alison Goate, Gerard Schellenberg, Daniel H. Geschwind, PSP genetics study group, John F. Crary, Adam Naj

**Author notes:** indicates corresponding authors contributed equally (John F. Crary, Adam Naj). The PSP genetics study group is a multisite collaboration including: German Center for Neurodegenerative Diseases (DZNE), Munich; Department of Neurology, LMU Hospital, Ludwig-Maximilians-Universität (LMU), Munich, Germany (Franziska Hopfner, Günter Höglinger); German Center for Neurodegenerative Diseases (DZNE), Munich; Center for Neuropathology and Prion Research, LMU Hospital, Ludwig-Maximilians-Universität (LMU), Munich, Germany (Sigrun Roeber, Jochen Herms); MRC Centre for Neurodegeneration Research, King’s College London, London, UK (Claire Troakes); Movement Disorders Unit, Neurology Department and Neurological Tissue Bank and Neurology Department, Hospital Clínic de Barcelona, University of Barcelona, Barcelona, Catalonia, Spain (Ellen Gelpi; Yaroslau Compta); Department of Neurology and Netherlands Brain Bank, Erasmus Medical Centre, Rotterdam, The Netherlands (John C. van Swieten); Division of Neurology, Royal University Hospital, University of Saskatchewan, Canada (Alex Rajput); Australian Brain Bank Network in collaboration with the Victorian Brain Bank Network, Australia (Fairlie Hinton), Department of Neurology, Hospital Ramón y Cajal, Madrid, Spain (Justo García de Yebenes).

## Abstract

Progressive supranuclear palsy (PSP) is a rare Parkinsonian disorder characterized by problems with movement, balance, cognition, and other symptoms. PSP differs from Alzheimer’s disease (AD) and other neurodegenerative diseases displaying abnormal forms of the microtubule-associated protein tau (“tauopathies”) by the presence of pathology not only in neurons, but also in astrocytes and oligodendrocytes. Genetic contributors may mediate these differences, however much of PSP genetics remains unexplained. Here we conducted the largest genome-wide association study (GWAS) of PSP to date including 2,779 cases (2,595 neuropathologically-confirmed) and 5,584 controls and identified six independent PSP susceptibility loci with genome-wide significant (*p* < 5×10^-8^) associations including five known (*MAPT*, *MOBP*, *STX6*, *RUNX2*, *SLCO1A2*) and one novel locus (*C4A*). Integration with cell type-specific epigenomic annotations revealed a unique oligodendrocytic signature that distinguishes PSP from AD and Parkinson’s disease. Candidate PSP risk gene prioritization using expression quantitative trait loci (eQTLs) identified oligodendrocyte-specific effects on gene expression in half of the genome-wide significant loci, as well as an association with elevated *C4A* expression in bulk brain tissue which may be driven by increased *C4A* copy number in PSP cases. Finally, histological studies demonstrated abnormal tau aggregates in oligodendrocytes that colocalize with C4 (complement) deposition. Integrating GWAS with functional studies including epigenomic and eQTL analyses, we identified potential causal roles for variation in *MOBP*, *STX6*, *RUNX2*, *SLCO1A2*, and *C4A* in the pathogenesis of PSP.

## Introduction

Tau proteinopathies (“tauopathies”), characterized by abnormal aggregates composed of the microtubule-associated protein tau inclusions, are a class of neurodegenerative diseases in aged individuals with varying yet overlapping clinical features including dementia, movement disorder, motor neuron disease, and psychiatric changes^1,2^. Among these is progressive supranuclear palsy (PSP; MIM #601104), a rare, late-onset neurodegenerative disease characterized by impaired movement, with symptoms including slowed movement (bradykinesia), loss of balance, frequent falls, and difficulty with eye movement (vertical supranuclear gaze palsy), as well as cognitive decline. Though an uncommon cause of dementia compared to AD, PSP is estimated to affect 5-17 per 100,000 persons in the US, making it the second leading cause of parkinsonism after PD, and autopsy studies have found PSP pathology in 2-6% of individuals with no PSP diagnosis prior to death, suggesting that it is more prevalent that appreciated in living individuals^3-5^. Given the sharing of common tau pathology across multiple neurodegenerative diseases, insights into the pathogenesis of PSP may yield potential therapeutic targets for a multitude of related diseases.

Key insights into PSP have come from decades of genetic studies which have demonstrated the disorder to be almost entirely sporadic disease, however careful clinical evaluation has revealed a tendency for family clustering^6,7^. While the chromosome 17q21.31 H1/H2 haplotype, an approximately 900Kb inversion polymorphism encompassing the gene encoding the tau protein, *MAPT*, remains the strongest known genetic risk factor for PSP (OR ≈ 5.5), loci containing variants with more modest effects have also been identified. These include variants associated with genome-wide significance (GWS; *P*<5×10^−8^) in or near genes *STX6*, *EIF2AK3*, and *MOBP*^8^. Subsequent work identified a variant, rs2242367, in an intron of the chromosome 12q12 gene *SLC2A13*, approximately ∼200kb upstream of the established PD/parkinsonism gene *LRRK2* as a locus influencing PSP disease duration^9^. Additional novel PSP risk loci were identified at chromosomes 6p12.1 (near *RUNX2*) and 12p12.1 (near *SLCO1A2*), as well as a unique 17q21.31 association in *MAPT*-adjacent gene *KANSL1^10,11^*. These loci help resolve only a portion of PSP heritability, and much of the genetic risk remains unexplained^10^.

Given that PSP is an archetypical primary tauopathy, advancing our understanding of the genetic architecture of the disorder has the potential to provide important insight into a number of diseases, but this requires large and robustly characterized cohorts^2,12^. Prior genome-wide association studies (GWAS) of PSP have been limited by relatively modest sample sizes, suboptimal control groups, and a paucity of downstream analyses to nominate causal genes surrounding or within the significant risk loci^8,10,11^. Here we performed the largest genetic association study of PSP to date, including 2,779 cases (of which 2,595 were autopsy-confirmed, building upon the 1,069 autopsy confirmed cases from stage 1 of work from 2011^8^) and 5,584 age-matched, non-demented, autopsy confirmed controls derived from the Alzheimer Disease Genetics Consortium^13^. We performed functional follow-up on each identified locus using a battery of annotation tools to pinpoint candidate causal genes and performed additional validation using molecular and histological approaches in human postmortem brain tissue. Taken together, our findings identify novel PSP genetic risk architechture and provide new functional insight into the genetic and molecular mechanisms driving tau proteostasis in PSP.

## Results

### Dataset collection and quality control

The cohort consisted of 2,779 PSP cases and 5,584 age-matched controls of which a majority were neuropathologically confirmed representing the largest PSP GWAS to date (**Table 1**). On average, the age-at-death of the controls was approximately ten years older than the cases and there were proportionally more females in the control population (60%) than in the cases (45%) To harmonize genotype data across multiple genotyping platforms and account for separate ascertainment of some sets of cases and controls, we constructed and implemented a genotype harmonization pipeline after quality control and prior to imputation (**Supplementary Figure 1**). After imputation of individual case-control sets to the TOPMed R5 reference panel, we filtered down to an overlapping set of 7,230,420 common (minor allele frequency (MAF)>0.01), high quality (Imputation *R^2^*>0.8) SNPs.

**Table 1.**
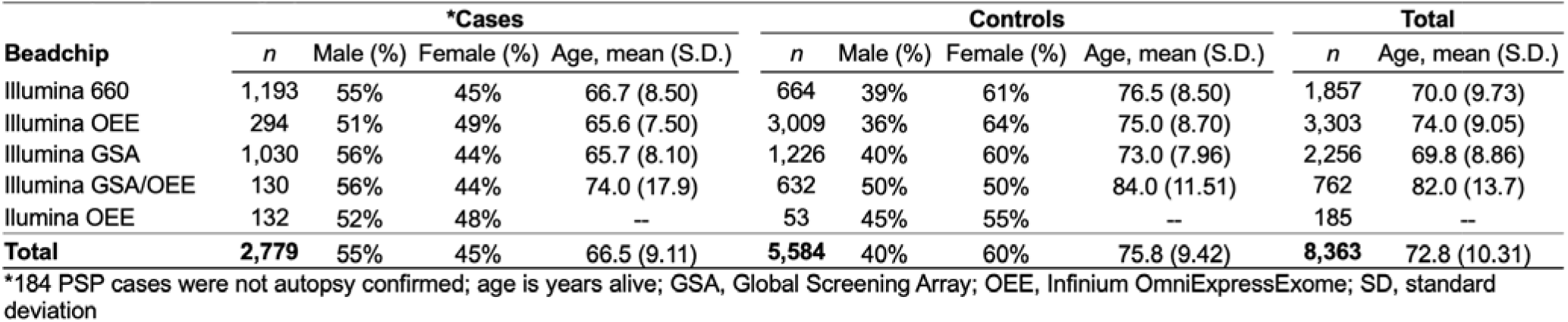
Subject demographics.

| Beadchip | *Cases |  |  |  | Controls |  |  |  | Total |  |
| --- | --- | --- | --- | --- | --- | --- | --- | --- | --- | --- |
|  | *n* | Male (%) | Female (%) | Age, mean (S.D.) | *n* | Male (%) | Female (%) | Age, mean (S.D.) | *n* | Age, mean (S.D.) |
| Illumina 660 | 1,193 | 55% | 45% | 66.7 (8.50) | 664 | 39% | 61% | 76.5 (8.50) | 1,857 | 70.0 (9.73) |
| Illumina OEE | 294 | 51% | 49% | 65.6 (7.50) | 3,009 | 36% | 64% | 75.0 (8.70) | 3,303 | 74.0 (9.05) |
| Illumina GSA | 1,030 | 56% | 44% | 65.7 (8.10) | 1,226 | 40% | 60% | 73.0 (7.96) | 2,256 | 69.8 (8.86) |
| Illumina GSA/OEE | 130 | 56% | 44% | 74.0 (17.9) | 632 | 50% | 50% | 84.0 (11.51) | 762 | 82.0 (13.7) |
| Illumina OEE | 132 | 52% | 48% | — | 53 | 45% | 55% | — | 185 | — |
| **Total** | **2,779** | **55%** | **45%** | **66.5 (9.11)** | **5,584** | **40%** | **60%** | **75.8 (9.42)** | **8,363** | **72.8 (10.31)** |

\*184 PSP cases were not autopsy confirmed; age is years alive; GSA, Global Screening Array; OEE, Infinium OmniExpressExome; SD, standard deviation

### Genome-wide association study

We observed six genome-wide significant loci (GWS; *P*<5×10^−8^), of which one was novel on 6p21.32 near *TNXB* at SNP rs369580 (OR [95% CI]: 1.43 [1.28, 1.60]; *P*=8.11×10^-10^) (**Table 2**). Our results confirm previously identified signals in loci containing *MAPT*, *STX6*, *MOBP*, *RUNX2*, and *SLCO1A2*. We observed a signal below the GWS threshold corresponding to the locus containing *EIF2AK3* (*P*=3.63×10^−5^) which was reported in previous studies^10^ (**Figure 1a**). Genome-wide associations demonstrated modest genomic inflation with λ=1.074 (**Supplementary Figure 2**). We performed stratified LD-score regression (S-LDSC), a method to estimate SNP heritability enrichment in sets of variants grouped by genomic features, using epigenomic annotations from four major CNS cell types assigning variants to promoters and enhancers^14-16^. We compared our S-LDSC results to recent GWAS conducted in Alzheimer disease (AD) and Parkinson’s disease (PD)^17-19^. As previously shown, AD heritability was enriched within microglial enhancers, whereas PD heritability was enriched in neuronal and oligodendrocyte promoters. In PSP, we observed a nominally significant enrichment in oligodendrocyte enhancer sequences (*P*=0.027) (**Figure 1b**). We then performed functional fine-mapping with PolyFun-FINEMAP, which estimated independent sets of variants (credible sets) at four loci, excluding the HLA-adjacent 6p21.32 (*TNXB*) and *MAPT* loci due to LD complexity^20,21^. At each locus we observed between one and three credible sets each containing 1-3 SNPs with a posterior inclusion probability (*PIP*)>0.1 (**Figure 1c**). We then overlapped the fine-mapped SNPs with the same epigenomic annotations as before, identifying several loci that contain fine-mapped SNPs overlapping CNS cell type epigenomic annotations. We also compared our fine-mapped SNPs to significant SNPs found in a recent massively parallel reporter assay (MPRA) using a previous PSP GWAS^22^. Only the 3p22.1 locus contained previously tested SNPs.

**Figure 1.**
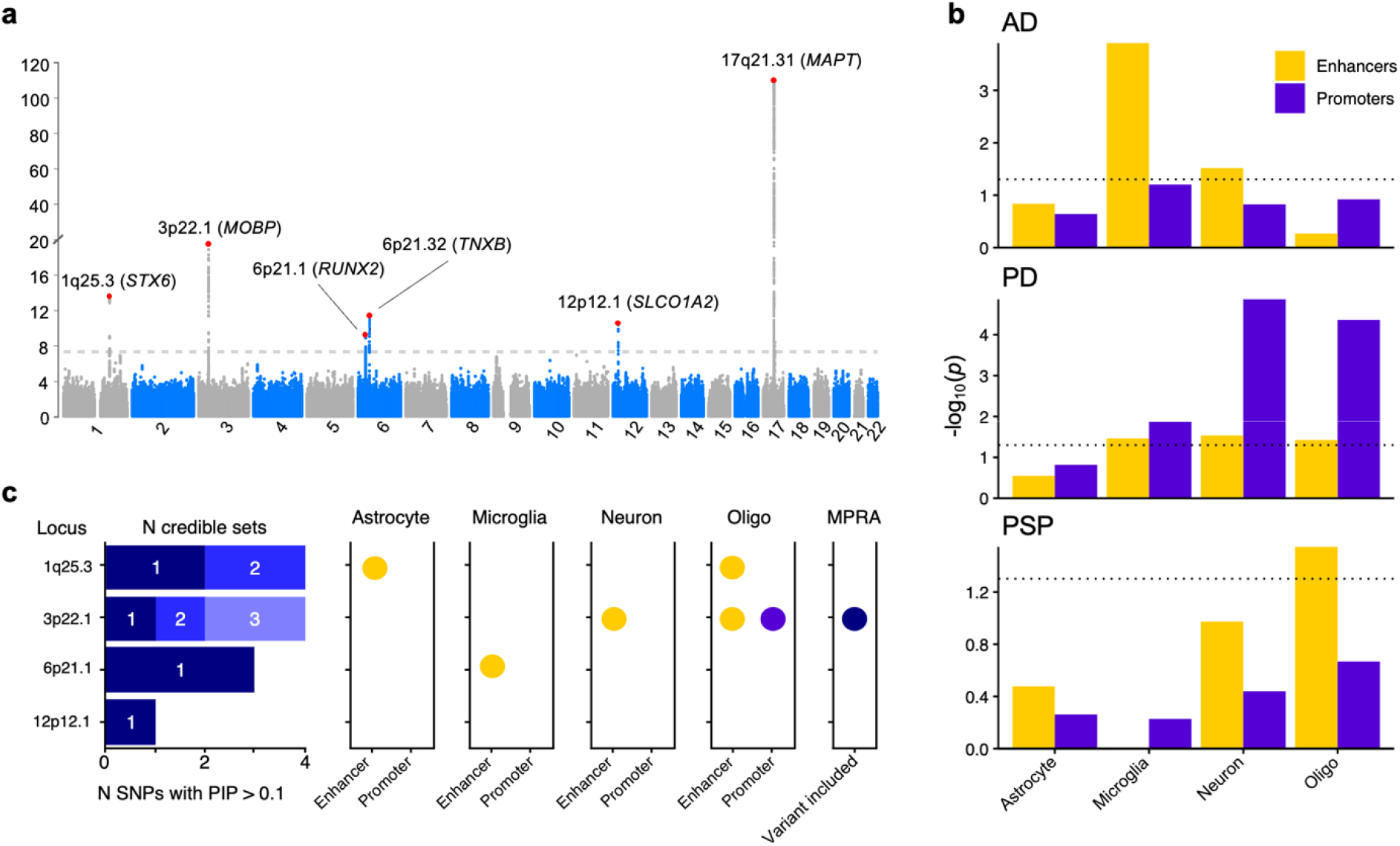
Common genetic variation in progressive supranuclear palsy (PSP) overlaps with cell type-specific epigenomic annotations. (a) Manhattan plot for PSP genome-wide association study (*n*=2,779 cases versus *n*=5,584 controls) identified six genome-wide significant loci (p<5 x 10^-8^) indicated by the dotted line containing approximately 218 genes around a 2-Mb window from the lead SNP. (b) LD-score regression using cell type-specific regulatory elements (dotted line indicates *p* < 0.05) derived from ChIP-seq^10^ demonstrates a unique cell-specific profile in PSP when compared to Alzheimer’s disease (AD) and Parkinson’s disease (PD)^18,19^. (c) Fine-mapping of the four amenable loci identified different numbers of causal SNPs and credible sets, some of which overlapped with cell type-specific regulatory elements and previously identified regulatory SNPs from an MPRA^22^. Oligo, oligodendrocytes; MPRA, Massively parallel reporter assay.

**Table 2.** Top SNPs.

| CHR | POS | Notation | rsID | Nearest Gene | Ref | Alt | FRQ A | FRQ U | R<sup>2</sup> | O.R. | S.E. | *p* |
| --- | --- | --- | --- | --- | --- | --- | --- | --- | --- | --- | --- | --- |
| 17 | 44101563 | 17q21.31 | rs9468 | *MAPT* | T | C | 0.942 | 0.7696 | 1.038 | 4.812 | 0.07 | $1.94 \times 10^{-110}$ |
| 1 | 180943529 | 1q25.3 | rs1044595 | *STX6* | C | T | 0.471 | 0.4064 | 0.989 | 1.347 | 0.039 | $2.92 \times 10^{-14}$ |
| 6 | 45454844 | 6p21.1 | rs12197948 | *RUNX2* | A | G | 0.705 | 0.644 | 1.008 | 1.328 | 0.041 | $5.43 \times 10^{-12}$ |
| 3 | 39508968 | 3p22.1 | rs631312 | *MOBP* | G | A | 0.362 | 0.2841 | 0.989 | 1.461 | 0.041 | $4.60 \times 10^{-20}$ |
| 12 | 21467215 | 12p12.1 | rs7966334 | *SLCO1A2* | C | G | 0.086 | 0.0546 | 0.967 | 1.664 | 0.077 | $3.66 \times 10^{-11}$ |
| 6 | 32020238 | 6p21.32 | rs369580 | *TNXB* | A | G | 0.886 | 0.8571 | 0.999 | 1.432 | 0.058 | $8.11 \times 10^{-10}$ |

ref, reference allele; Alt, alternate allele; FRQ A, allele frequency in cases; FRQ U, allele frequency in controls; R<sup>2</sup>, quality; O.R., Odds ratio; S.E., Standard error.

### Colocalization analysis

We then sought to identify specific causal genes at each locus by incorporating expression quantitative trait loci (eQTLs) from bulk brain expression data from the Genotype Tissue Expression (GTEx) project and expression data from sorted CNS cell types from single nucleus RNA-seq (snRNA-seq)^23^ (**Figure 2**). We performed colocalization tests estimating the probability that the same single causal variant is associated with both disease risk and with gene expression by comparing all matching SNPs within each GWAS locus (±1Mbp) to those tested in each eQTL. This approach prioritized *STX6* and *RUNX2* as the likely causal genes in the 1p25.3 and 6p21.1 loci respectively as they had a high posterior probability (*PP4*) across multiple brain regions. In the complex HLA-adjacent 6p21.32 locus, *BTNL2* colocalized in only three brain regions. Cell type-specific eQTLs identified *STX6* and *RUNX2* in oligodendrocytes at *PP4*>0.8, whereas eQTLs for *MOBP* were found in both oligodendrocytes and excitatory neurons^23^. No gene was prioritized by eQTLs in the 12p12.1 locus. In summary, three of the six GWAS loci have evidence of acting through gene expression in oligodendrocytes. Guided by these cell type-specific colocalizations, we examined our loci more closely.

**Figure 2.**
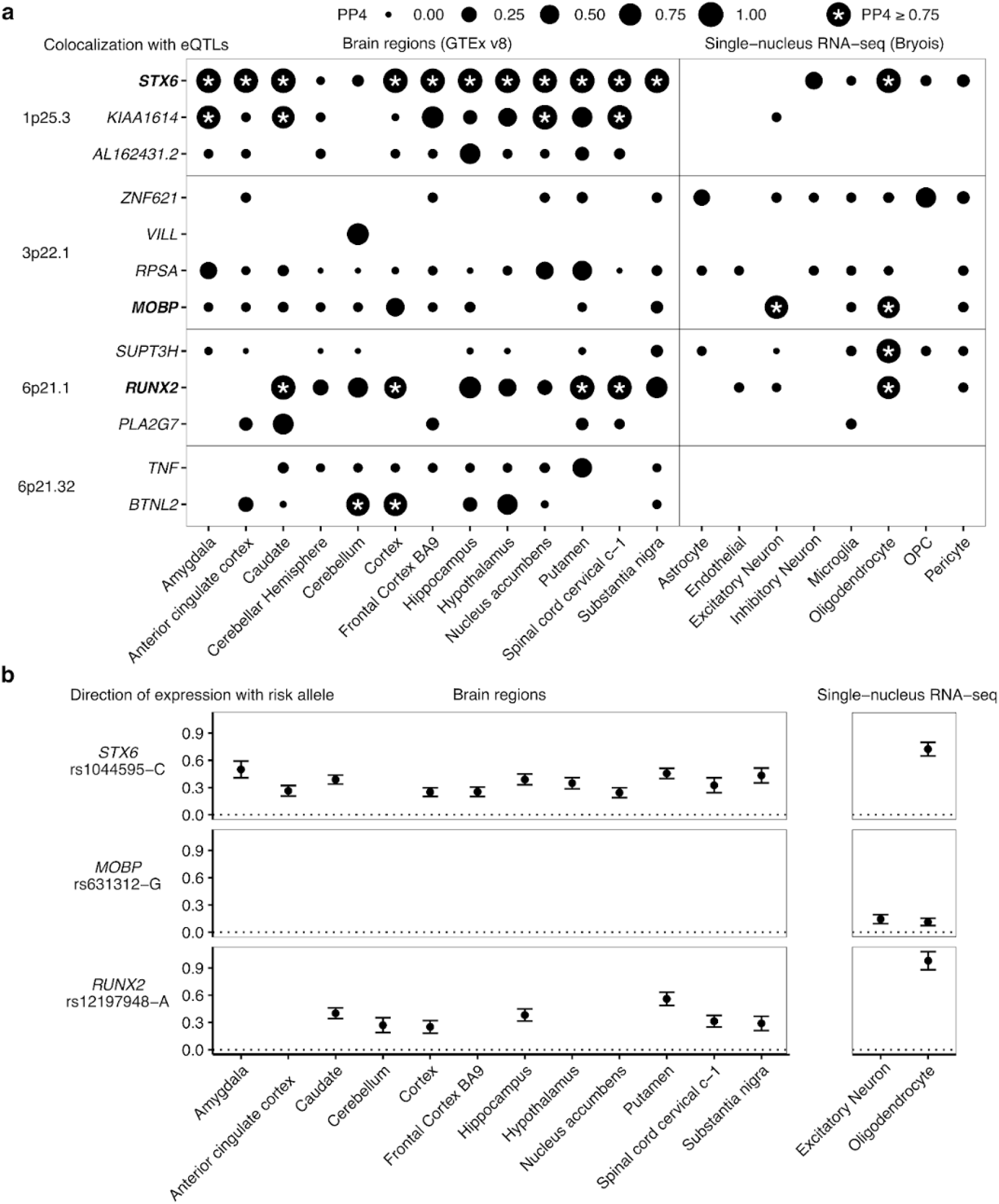
Progressive supranuclear palsy (PSP) candidate risk gene prioritization. Datasets were GTEx (bulk RNA-seq) and Bryois et al (single nucleus RNA-seq) eQTLs across brain regions and purified cell types analyzed with COLOC. a) All genes and loci shown with PP4L>L0.5 in at least one eQTL dataset. Point sizes are scaled to the magnitude of posterior probability of colocalization (PP4). * PP4 >= 0.75. OPC, Oligodendrocyte precursor cell. b) Directions of effect on gene expression for *STX6*, *MOBP*, and *RUNX2* relative to the risk allele (increased in frequency in PSP cases) of the lead GWAS SNP. Error bars denote standard error of the eQTL association.

*1p25.3 - STX6.* Colocalization of 1p25.3 prioritized *STX6* in multiple brain regions and with oligodendrocyte-specific eQTLs. The risk allele of the lead GWAS SNP rs1044595-C is associated with increased expression of *STX6* in bulk brain samples and in purified oligodendrocytes (**Figure 2b).** Fine-mapping of 1p25.3 identified two credible sets each containing 2 SNPs with a posterior inclusion probability > 0.1. These SNPs overlapped with oligodendrocyte enhancer sequences as identified by cell type-specific ChIP-seq (**Figure 3a**). The first credible set included the lead GWAS SNP rs1044595 as well as a second SNP rs3789362, in high LD (R^2^=0.96, 1000 Genomes European superpopulation). rs2789362 overlaps with an oligodendrocyte-specific enhancer within an intron of *STX6*. Taken together, this suggests that rs3789362-A increases *STX6* expression by modifying an oligodendrocyte-specific enhancer sequence.

**Figure 3.**
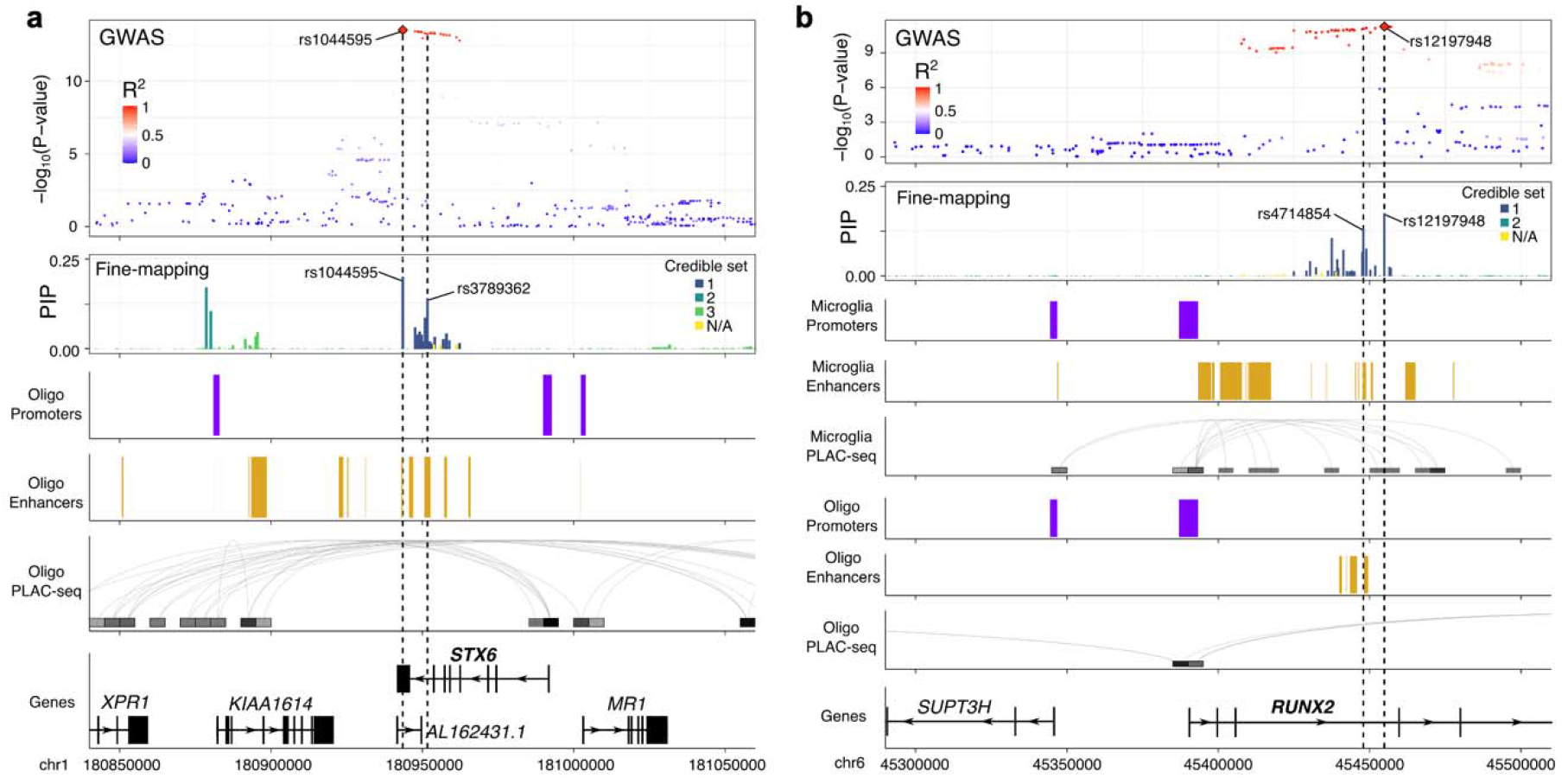
Fine-mapping and cell type-specific epigenetic features in the *STX6* and *RUNX2* loci. Functional annotations from cell type-specific regulatory regions (enhancers and promoters) and cell type-specific DNA interactome anchors from proximity ligation-assisted ChIP-Seq (PLAC-seq) are shown in the loci containing *STX6* (a) and *RUNX2* (b). SNPs are colored by their LD with the lead GWAS SNP in Europeans (1000 Genomes European superpopulation).

*3p22.1 - MOBP*. As a well-established cell-type marker for oligodendrocytes, we expected to see strong colocalization between the 3p22.1 locus and *MOBP* in bulk brain samples and oligodendrocytes specifically. The risk allele of the lead GWAS SNP rs631312-G is associated with increased *MOBP* expression in oligodendrocytes (**Supplementary Figure 3**). We observed three single-variant credible sets in 3p22.1, two of which overlapped with a region at the transcription start site of the *MOBP* gene containing ChIP-seq regions defined as both promoters and enhancers in oligodendrocytes and densely connected by proximity ligation-assisted ChIP-seq (PLAC-seq) contacts (**Supplementary Figure 3**)*^16^*. Taken together, this suggests that variants within 3p22.1 alter *MOBP* expression in oligodendrocytes specifically.

*6p21.1 - RUNX2.* Colocalization of 6p21.1 prioritized *RUNX2* in multiple brain regions as well as in oligodendrocytes specifically, with the risk allele of the lead GWAS SNP rs12197948- A associated with increased levels of *RUNX2* expression in all datasets (**Figure 2b**). Fine-mapping identified a single credible set containing 3 SNPs with a PIP>0.1. Two of the SNPs (rs12197948 and rs4714854, R^2^=0.99, 1000 Genomes European superpopulation) sit within the third intron of *RUNX2*, which contains an annotated enhancer in both microglia and oligodendrocytes (**Figure 3b**). Although rs4714854 overlaps a microglia enhancer peak, it is also very close to the oligodendrocyte enhancer peak. Additionally, PLAC-Seq in microglia identified several contacts between the microglia enhancer region and the *RUNX2* promoter. Therefore, although colocalization suggests oligodendrocytes to be the causal cell-type, we cannot rule out that the GWAS association at 6p21.1 may also affect *RUNX2* expression in microglia through a shared enhancer sequence.

*6p21.32 - C4A.* Although we observed colocalization with bulk brain eQTLs for *BTNL2*, we reasoned that the known LD complexity in the locus may obscure genuine colocalizations. As an alternative approach we applied INFERNO, which uses LD pruning at each GWAS locus to construct a set of independent SNP sets, which can then be tested for eQTL colocalization separately, which is the main difference COLOC and INFERNO^24,25^. In the 6p21.32 locus, INFERNO identified 3 sets of SNPs (**Figure 4a**). Each SNP set was then colocalized with all nearby genes (**Figure 4b**) using eQTLs from 13 GTEx brain regions (v7). We observed multiple genes in the locus to be colocalized, with the largest number of colocalizations (*PP4*>0.9) in all 3 SNP sets being with eQTLs for the gene *C4A* (**Figure 4c**). Comparing each the *P*-value of association with PSP (*P*_GWAS_) with the eQTL association in GTEx Frontal Cortex (*P*_eQTL_), we observed that while including all SNPs from the locus resulted in a minimal probability of colocalization (*PP4*=2×10^-5^; **Figure 4d**), using the individual sets of SNPs resulted in a much higher colocalization for all 3 sets with *C4A* eQTLs (*PP4*>0.8; **Figure 4e**). The risk allele of the lead GWAS SNP rs2523524-G was associated with increased *C4A* expression (**Supplementary Figure 4**). Given the known C4A copy number variation in this locus we hypothesized that variability in *C4A* copy numbers could explain the observed signal. To test this hypothesis, we generated imputed *C4A* and *C4B* copy number values and ran logistic regression for case control status based on alterations in the copy number of each gene and observed that *C4A* copy number was nominally significantly associated with PSP status (*p* = 1.58 × 10^-6^), whereas *C4B* copy number was not (*p* = 0.14). Additionally, we ran our association study including either *C4A* or *C4B* copy number status as a covariate and observed that the inclusion of only *C4A* copy number status was able to reduce the signal observed in the locus below the genome-wide threshold (**Figure 5a-c).** We therefore nominated *C4A* as the most likely causal gene at this locus.

**Figure 4.**
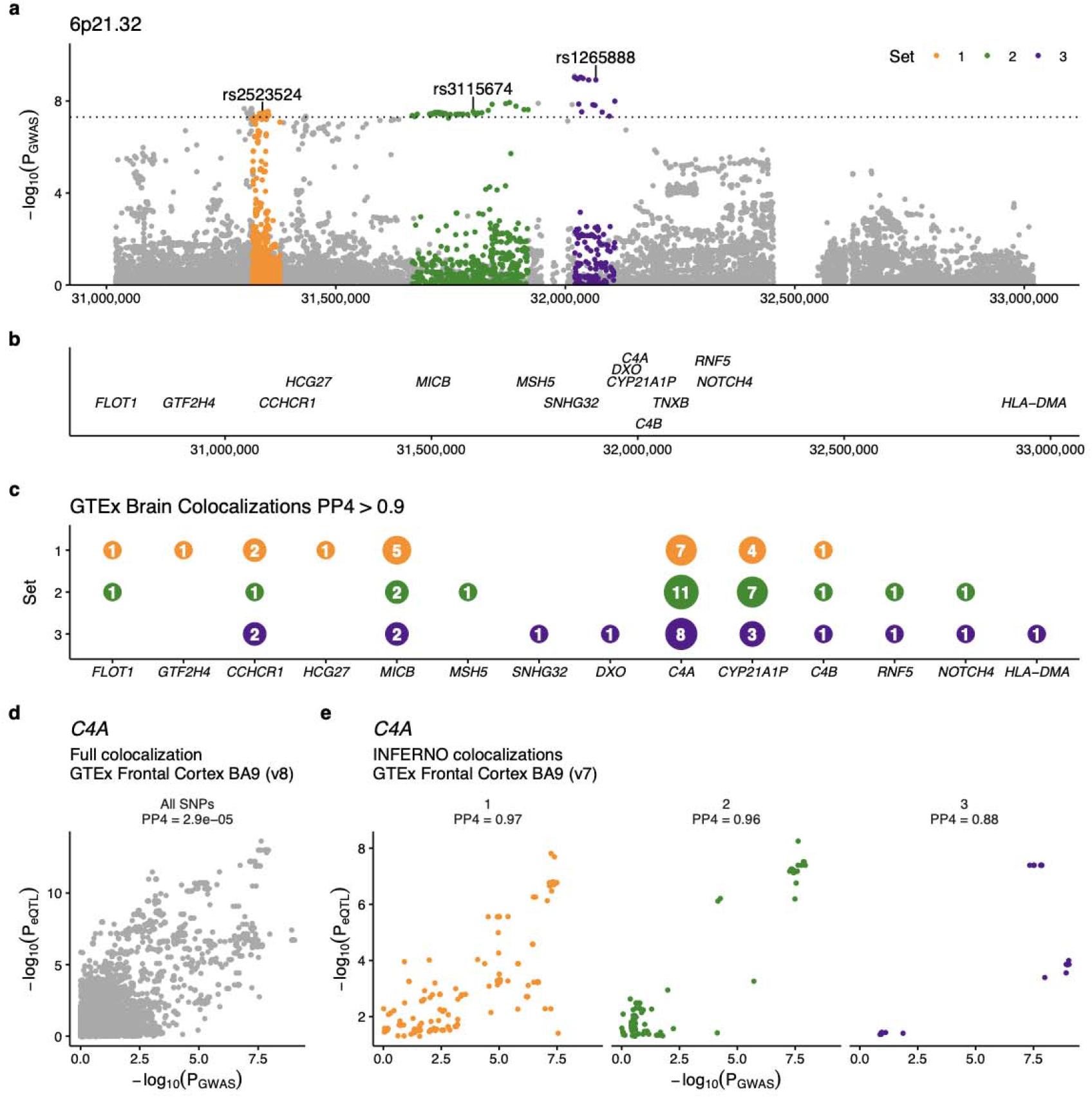
Risk gene prioritization implicates *C4A* as a candidate gene at 6p21.32. (a) LD pruning selects three sets of credible SNPs in the locus which surround several genes (b). Colocalization of each SNP set with expression quantitative trait loci in multiple brain regions (GTEx v7) (c) shows that an eQTL for *C4A* is present in the majority of tested brain regions, more than any other gene, with a high posterior probability of colocalization (PP4) > 0.9. (d) Comparing the relationship between the P-values found in GWAS (P_GWAS_) and the *C4A* eQTL (P_eQTL_). When using all SNPs in the locus there is a low PP4 with *C4A* eQTLs in the frontal cortex (PP4 < 0.01), whereas the three sets of SNPs chosen by INFERNO all show high probability of colocalization (PP4 > 0.8).

**Figure 5.**
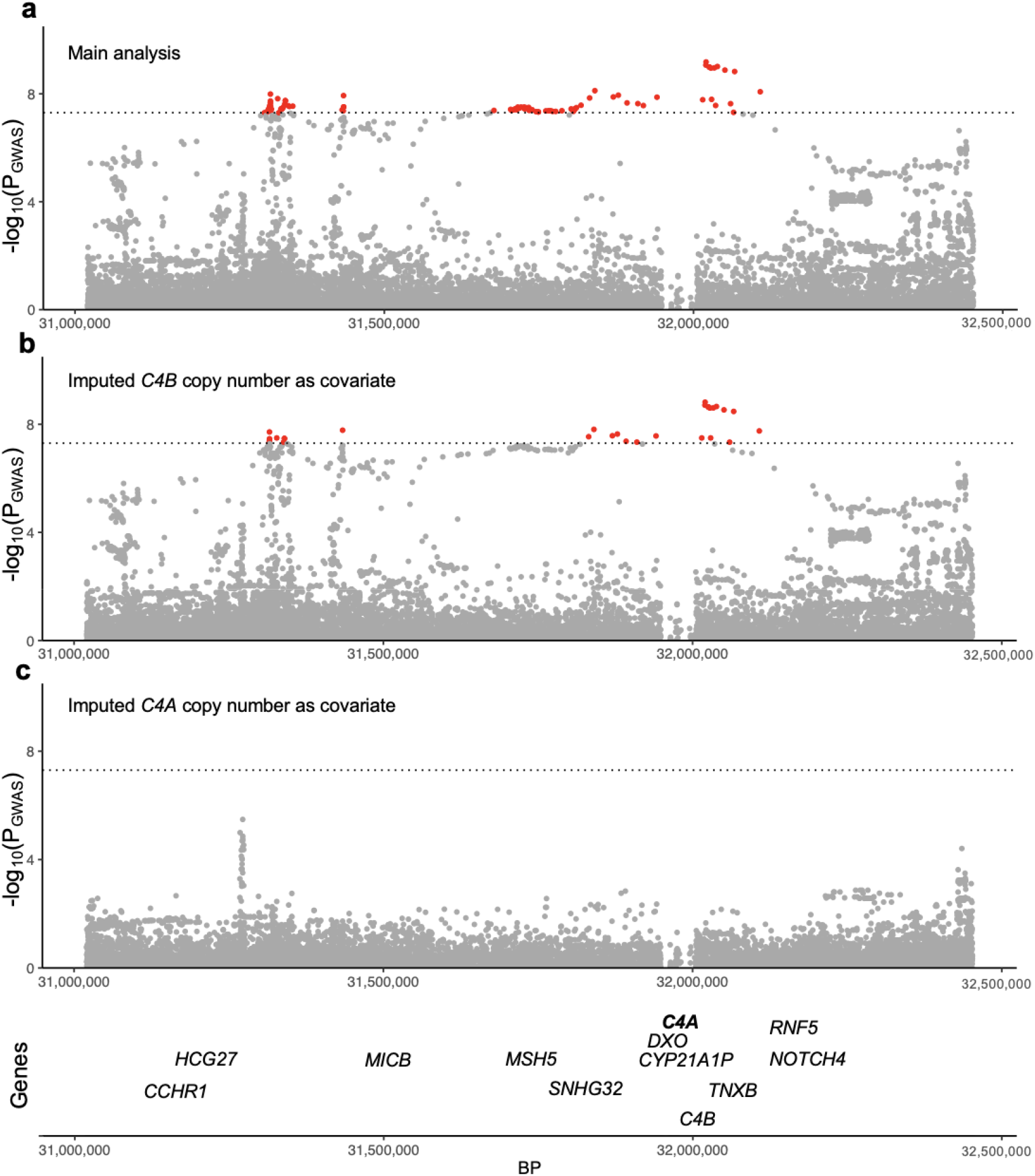
The addition of imputed C4A copy number to the association model reduces the observed signal below the genome-wide threshold suggesting its role as the candidate gene in the loci. (a) Despite losing 14 cases due to the inability to impute C4A and C4B copy number, a genome-wide significant signal is still observed in 6p21.32. (b) The addition of imputed C4B copy number to the association model still produces a robust genome-wide significant signal, whereas adding imputed C4A copy number (c) pushes the signal below the genome-wide threshold suggesting that imputed C4A copy number is influencing the signal.

### Differential gene expression in frontal cortex and cerebellum

Given the genetic, eQTL, and fine-mapping evidence suggesting that multiple genes contained in the GWS-associated loci are potentially implicated in PSP, we leveraged a previously generated bulk RNA-seq dataset to identify regionally specific changes in gene expression in the frontal cortex and cerebellum in patients with PSP compared to non-neurological disease controls^26^. After re-analyzing the raw data to include relevant covariates, we focused only on the 16 candidate genes identified in the significant GWAS loci. In the frontal cortex, we observed significantly increased expression of *STX6* and *FLOT1*, and significantly decreased expression of *PLA2G7*, *MOBP*, *MSH5*, *HLA-DPB1*, *HLA-DMB*, and *SLCO1A2* in PSP versus controls (*P*<0.0025, **Figure 6a**). In the cerebellum, the same significantly decreased expression pattern was observed in *MOBP*, *MSH5*, and *SLCO1A2*, but no significant differences were observed in *HLA-DPB1*, *HLA-DMB*, and *FLOT1* (*P*>0.0025, **Figure 6b**). The data suggest regionally specific changes of multiple genes identified in loci identified from the GWAS data which may have downstream effects on disease relevant protein expression.

**Figure 6.**
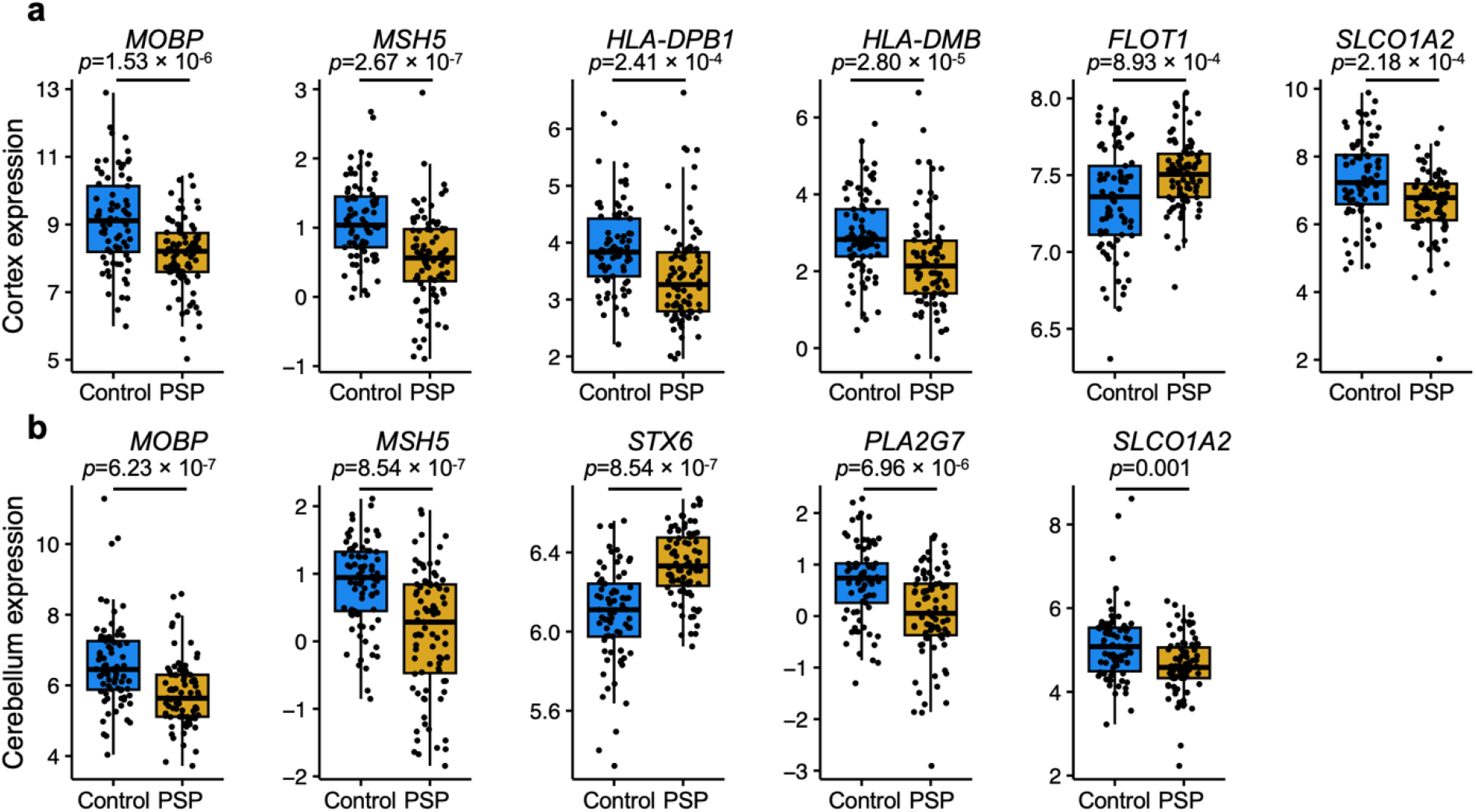
Differential expression of progressive supranuclear palsy (PSP) candidate risk genes. Bulk RNA-seq from PSP (*n*=84 cortices, 83 cerebellums) and control (*n*=77 cortices, 77 cerebellums) human post-mortem brain tissue showing significant differences in (a) six neocortical genes and (b) five cerebellar. Expression differences shown are normalized read counts. Sixteen total candidate genes were present in the dataset. Bulk RNA-seq fold change values shown only if the differential expression p-value < 0.0025.

### Immunohistochemical and biochemical analysis

Given the novel genetic association observed in the 6p21.32 locus and the downstream computational evidence nominating *C4A* as a candidate gene, we examined tissues from both PSP cases and controls biochemically and immunohistochemically to see if there was any cell type-specific pathology relevant to the *C4A* signal. As expected, multiplex immunohistochemical staining of controls (*n*=10) showed little hyperphosphorylated tau (p-tau, AT8) pathology and a minimal amount of C4A protein signal in the frontal cortex alongside positively stained healthy oligodendrocytes (OLIG2) (**Figure 7a, Supplementary Figure 5**). In PSP cases (*n*=10), we observed a strong immunohistochemical p-tau signal in neurons and tufted astrocytes, a hallmark pathology of the disorder, as well as a marked increase in C4A protein expression in the axons in association with p-tau positive oligodendrocytic coiled bodies (**Figure 7b, Supplementary Figure 5**). Image analysis revealed significantly more C4A staining in PSP than controls (*p*=0.0001, **Figure 7c**) Furthermore, these axonal profiles were abnormal, being significantly shorter in PSP than controls (*p*=0.001, **Figure 7d**). Finally, quantitative immunoblots of the C4A alpha chain showed significantly higher levels in PSP (*n*=6) compared to controls (*n*=7, *P*=0.008, **Figure 7e, Supplementary figure 6**). Together, these findings provide histological and biochemical evidence of C4 abnormalities in association with tau pathology in PSP brains.

**Figure 7.**
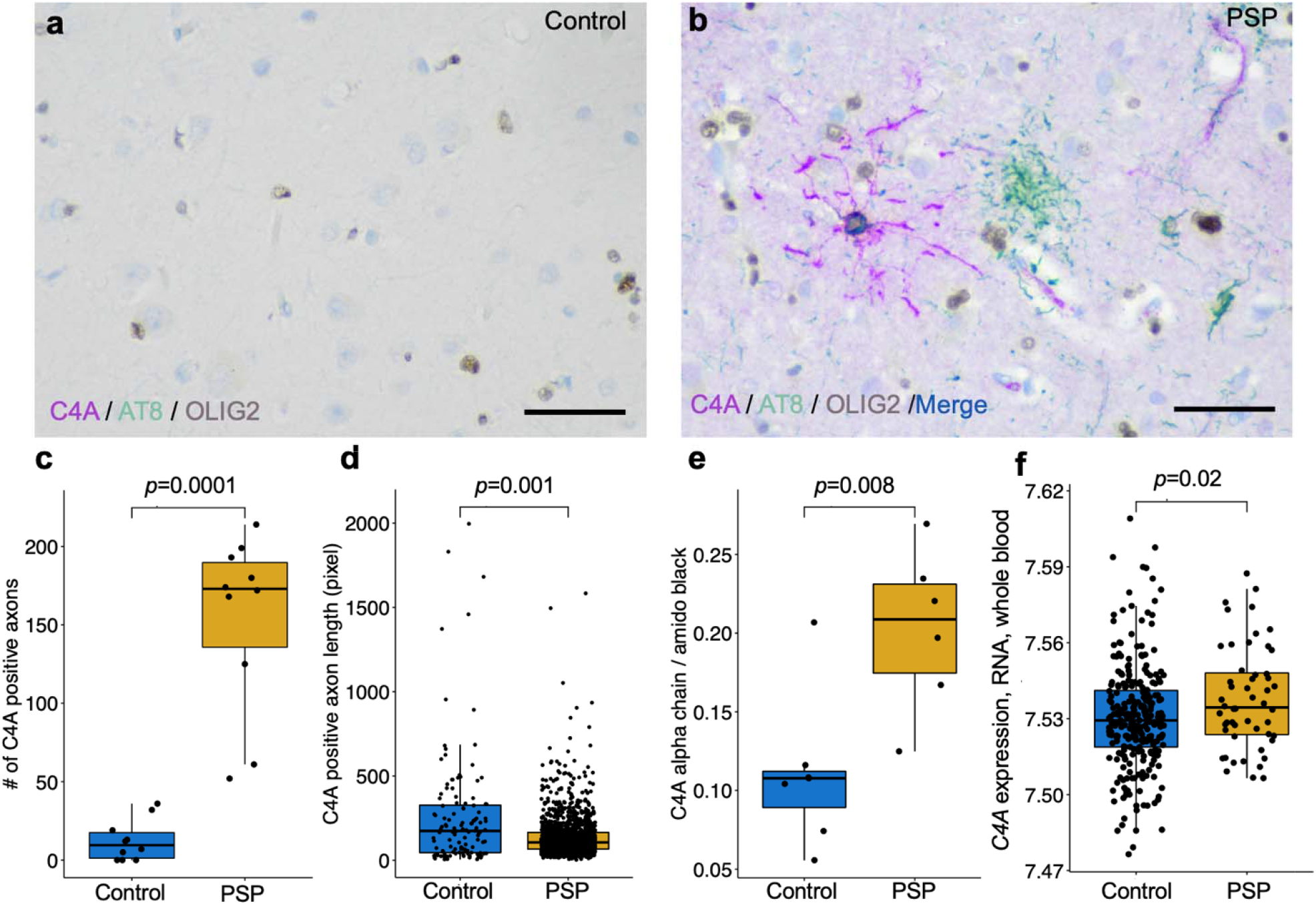
Elevation of C4A protein in the frontal cortex of human postmortem progressive supranuclear palsy (PSP) brain tissue. (a). Triple immunohistochemical stains of C4A (magenta), hyperphosphorylated tau (AT8, green), and OLIG2 (brown) show limited C4A and hyperphosphorylated tau staining in controls (b). Staining in PSP demonstrates colocalization of C4A and hyperphosphorylated tau in an OLIG2 positive coiled body (blue and brown). Tufted astrocyte cell bodies (green) and neurofibrillary tangles (not shown) are negative for C4A in PSP. The average number of C4A positive axons per subject (c) was significantly higher in PSP than controls (*n*=10 cases and controls, frontal cortex, 5 regions of interest per subject, 1,538 observations found in PSP, 124 in controls). Quantification of C4A positive axons revealed significantly shorter (d) axonal length in PSP when compared to controls. (e) Quantitative immunoblots of the C4A alpha chain showed significantly higher levels in PSP compared to controls (n=7 cases, 6 controls). (f) *C4A* gene expression in whole blood from patients with PSP was significantly higher than controls (n=51 cases, 281 controls). Scale bar is 50 micrometers.

### C4A expression in whole blood

As we observed elevated C4A protein in PSP oligodendrocytes postmortem, we reasoned that *C4A* mRNA may be elevated in living patient blood samples. We re-analyzed a publicly available RNA microarray dataset generated from whole blood from non-neurological controls (*n*=281) and clinically diagnosed PSP cases (*n*=51)^27^. We observed *C4A* mRNA to be upregulated in blood from PSP patients compared to controls (*P*=0.02, **Figure 7f, Supplementary Figure 7**).

## Discussion

We have summarized our ensemble of downstream genetic analyses in a single table (**Table 3**). In the 1q25.2 locus, we nominate *STX6* as the causal gene, as it shows eQTL colocalization in both bulk brain and in oligodendrocytes; the fine-mapped SNPs overlap an oligodendrocyte-specific enhancer region; and *STX6* gene expression is upregulated in the cerebellum of PSP brains. In the 3p22.1 locus, we nominate *MOBP* as the causal gene, although it does not show eQTL colocalization in bulk brain and others have suggested its role in *SLC25A38/Appoptosin* gene locus 70 kb away, we did observe a signal in oligodendrocyte single cell data as well as downregulation of expression in PSP brains in addition to fine mapping oligodendrocytic enhancers and promoters as well as an observed signal in the MPRA data^22,28^. In the 6p21.1 locus we nominate *RUNX2* based on similar observations however there is some discrepancy over the cell type-specificity given the signal mapped to microglial enhancers but also had an oligodendrocytic eQTL signal. We did not observe a signal in the loci containing *EIF2AK3* and *LRRK2* which have been previously observed, albeit *LRRK2* was found to be associated with PSP survival not susceptiablity^8,9^. In the novel 6p21.32 locus reported here, nomination of a causal gene was challenging given the limited fine mapping and eQTL interactions reported. Thus, we turned to biochemical and immunohistochemical validation which strongly supports *C4A*’s role in complement activation and neuroinflammation in PSP and thus calls for further exploration of the mechanistic role of this gene in PSP. Lastly, in the 12p12.1 locus we nominate *SLCO1A2* given the evidence provided for its significant downregulation of expression in PSP brains in multiple regions.

**Table 3.**
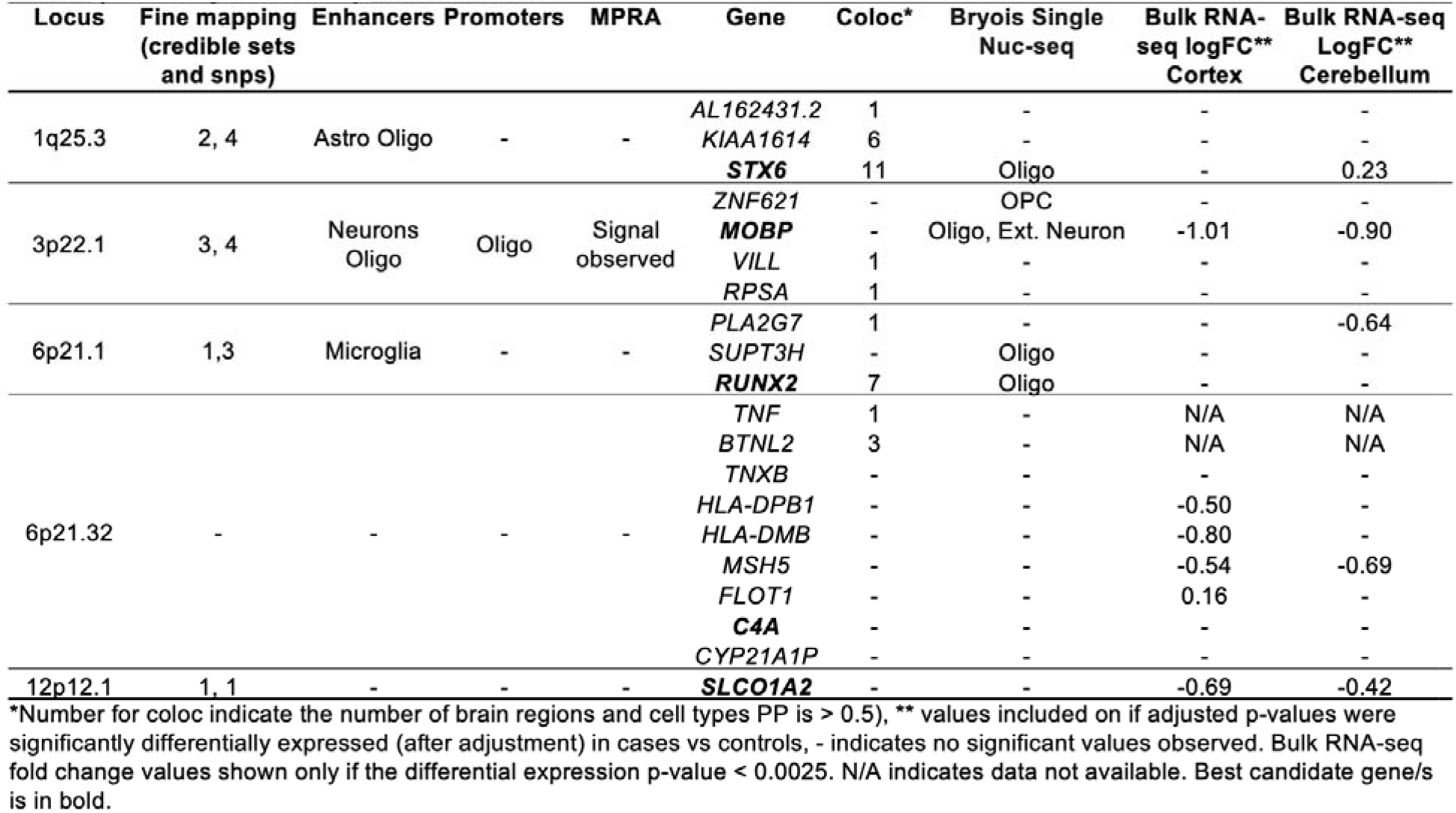
Summary of the computational results.

Tau proteinopathies, especially primary tauopathies such as PSP that arise independently of amyloid-β, hold significant clinical and scientific importance due to their prevalence and potential to provide novel mechanistic insights into neurodegeneration^2,12,29^. Furthermore, understanding abnormalities in tau dysfunction offers promise in advancing our knowledge of neurodegenerative diseases more broadly^30^. Tau-related neurodegeneration has been proposed to occur through various mechanisms, including apoptosis, excitotoxicity, oxidative stress, inflammation, mitochondrial dysfunction, prion-like propagation, and protein aggregation. However, the connection between these mechanisms and genetic drivers remains poorly understood^31^. Furthermore, the vulnerability of distinct cell populations in tauopathies remains incompletely characterized. Human GWAS continue to be an invaluable tool in elucidating causal mechanisms in neurodegenerative diseases^32^. Notably, increasingly large genetic studies of Alzheimer disease (AD) have continued to enable the discovery of new risk loci, including those related to neuroimmune meachanisms and other functions^32^. Genetic studies of the primary tauopathies, including PSP, have remained small despite their potential to highlight non-amyloid-driven mechanisms that are becoming increasingly relevant given the growing emphasis on combination therapy in AD^33^. This study of PSP, the prototype non-AD primary tauopathy, that includes 8,363 total subjects in a genome-wide study, sheds new light on these candidate mechanisms.

Our study uncovered a novel genetic signal at *C4A* which encodes the acidic form of complement factor 4, part of the classical activation pathway. This finding was further supported by histological, biochemical, and blood biomarker analyses. Critically, co-localization of C4A protein with abnormal tau species in oligodendrocytes further supports that innate immune function plays a causal role in driving this pathological interaction in PSP. *C4A* has also been identified in a GWAS of ALS, which is hypothesized to be a distal (dying-back) axonopathy, suggesting the possibility that axon-myelin interactions might contribute to and link these disorders^34,35^. Furthermore, there is a genetic link between copy number variation in *C4A* and schizophrenia and functional studies have shown overproduction of this protein promotes excessive synaptic loss and behavioral changes in mice^36-38^. To this point, we also observed this link when we ran our association study with imputed *C4A* copy number as a covariate resulting in reduction of our primary signal in 6p21.32 below the genome-wide threshold. Additionally, using a CRISPR deletion assay targeting a SNP contained within 6p21.32 (previously found in a GWAS of AD) reduced *C4A* expression in iPSC-derived astrocytes providing more evidence for the role of *C4A* in neurodegeneration^22^. Similar to what we observed in a whole blood dataset, others have observed that differences in complement protein levels in the cerebrospinal fluid across various neurodegenerative diseases^39-41^. Lastly, although it has been hypothesized for some time that complement activation is involved in neurodegeneration and this has been shown in murine models, our histopathological evidence in human post-mortem tissue shows marked morphological features in oligodendrocytes with p-tau pathology, demonstrating a link back to the identified novel genetic loci^38,42^. In summary, these findings underscore the importance of exploring the role of innate immune interactions and oligodendrocyte pathology in the pathogenesis of multiple neurodegenerative conditions.

Despite the insights gained from this study, several limitations should be considered when interpreting the findings. Although most cases were autopsy-confirmed to assure correct classification, this limited our overall sample size, compared to using clinically diagnosed, or even proxy cases. Firstly, the small sample size used limits the statistical power to detect small effect size loci. GWAS typically require very large sample sizes to achieve sufficient statistical power, and inadequate sample sizes can result in false-negative and false-positive findings, potentially missing true genetic associations. While we were able to provide biochemical and histological evidence prioritizing *C4A* at the 6p21.32 locus, this gene resides in the *HLA* region where complex structural genomic rearrangements complicate identification of causal variants. This also limits our ability to nominate genes at 17q21.31, thus there is a critical need for advanced computational tools and long-read sequencing in these loci. Follow up studies, such as functional genomic analyses, model organism experiments, and a replication cohort are necessary to elucidate how the identified variants affect biological processes related to neurodegeneration.

We explored the genetic risk for PSP, confirming previous signals while also identifying one novel association. Among the confirmed genetic signals, *MAPT* remains the strongest, consistent with the well-established role of the *MAPT* haplotypes in tau proteinopathies^43^. We also confirmed the association with myelin-associated oligodendrocytic basic protein (*MOBP)* and assigned this signal to gene expression in oligodendrocytes, in line with its role in synthesis and maintenance of myelin. *MOBP* is also a candidate risk gene in amyotrophic lateral sclerosis (ALS) and a previous colocalization analysis has shown that the same causal SNPs are found in PSP, ALS, and corticobasal degeneration^8,34,44,45^. *STX6* encodes syntaxin 6, a soluble N-ethylmaleimide-sensitive factor attachment protein receptor (SNARE) that localizes to endosomal transport vesicles, and has a critical role in intracellullar trafficking. Intriguingly, recent studies have implicated *STX6* in regulation of immune function, but it is also highly expressed in other cells including oligodendrocytes, which we also observed here in this study^46-48^. However, our investigation did not support the involvement of the *EIF2A* locus identified in previous PSP GWAS, even though parallel human tissue research has implicated the integrated stress response in tauopathy^49,50^, although this is controversial^51^. Thus, additional validation is warranted. We observed a signal in *RUNX2*, which encodes runt-related transcription factor-2, which had previously been identified but not replicated until this study^10,11^. Although we observed an association in oligodendrocytes, *RUNX2* is highly expressed in microglia and may play a role in regulation of phagocytosis^52-54^. Finally, we also replicated the signal at the *SLCO1A2* locus, which encodes the solute carrier organic anion transporter family 1A2 protein, which is also highly expressed in human oligodendrocytes, although we did not observe a colocalizing eQTL, suggesting that the locus may act through an alternative molecular mechanism^55^. *SLCO1A2* has been linked to beta-amyloid burden in AD suggesting a generalized role in brain homeostasis amongst tau proteinopathies^56^. Taken together, these findings highlight the potential significance of immune-related mechanisms in PSP and for the first time in the field, we have made use of cell type-specific data to nominate increased gene expression specifically in oligodendrocytes as the mechanism behind 3 out of 6 risk loci.

In conclusion, this manuscript presents findings from a GWAS of PSP, a neurodegenerative disease characterized by movement, cognitive, and behavioral impairments. The study identified six independent susceptibility loci associated with PSP, including a novel locus at 6p21.32. Through computational analyses and functional fine-mapping, several candidate genes were nominated, including *MOBP*, *STX6*, *RUNX2*, *SLCO1A2*, and *C4A*. Additionally, the study revealed a unique oligodendrocyte signature that distinguishes PSP from other neurodegenerative diseases. Further investigation of the identified susceptibility loci and their functional consequences, as well as the examination of cell specific pathologies, provides new insights into the genetic and molecular mechanisms underlying PSP. These findings contribute to our understanding of tau proteostasis and may have implications for related tauopathies.

## Methods

### Datasets

For a majority of the cases included in the study, inclusion criteria were a neuropathological diagnosis of PSP (*n*=2,595), with the exception of a small number of cases, both living and deceased, that only had a neurological diagnosis (*n*=184). PSP subjects with comorbid pathological features of other neurodegenerative disorders were not excluded from the study including AD-like features, Lewy bodies, and TDP-43 as prevalence of these comorbid features has been previously demostrated^57^. The controls had no clinical evidence of cognitive impairment or a movement disorder (*n*=5,584) and neuropathologically could only have age-related pathological changes. A full list of the institutions where the material was collected can be found in **Supplementary Table 1** and it should be noted many of the samples included here were contained in previous studies^8,10,11^.

### Genotyping and quality control

PSP cases and controls were genotyped at three different institutions (University of Pennsylvania, Icahn School of Medicine at Mount Sinai, and the University of California Los Angeles) on three genotyping platforms (Illumina Human660W, Illumina OmniExpress 2.5, and Illumina Global Screening Array) in 10 total batches (**Supplementary Figure 1**). DNA was isolated from the subjects’ using methods detailed elsewhere^8,11,58^. The cases and controls were genotyped at each of the respective institutions, merged, and harmonized to contain the same variants and single nucleotide polymorphism (SNP) and sample level quality control (QC, detailed below) was performed followed by imputation. The process was repeated by combining the data from the three centers and the overlapping variants were again harmonized. PLINK v1.9 was used to perform quality control. SNP exclusion criteria included minor allele frequency < 1%, genotyping call-rate filter less than 95%, and Hardy–Weinberg threshold of 1 × 10^−6^. Individuals with discordant sex, non-European ancestry, genotyping failure of > 5%, or relatedness of > 0.1 were excluded. A principal component analysis (PCA) was performed to identify population substructure using EIGENSTRAT v6.1.4 and the 1000 genomes reference panel. Samples were excluded if they were six standard deviations away from the European population cluster.

#### TOPMed imputation and post-processing

Each dataset was imputed on the Trans-Omics for Precision Medicine (TOPMed) Imputation Server (TIS) using the multi-ancestry release 5 (R5) reference panel which includes data on from 97,256 participants with 308,107,085 SNPs observed on 194,512 haplotypes^59,60^. Phasing was performed on the TIS using EAGLE with subsequent imputation using Minimac4^61,62^. Imputed variants were filtered using a conservative quality threshold, *R^2^*≥0.8, to assure high quality of variants, and additional filtering on variants overlapping all genotype sets with MAF>0.01 was performed prior to analysis.

#### Association analysis

Single-variant genome-wide association analyses was performed jointly on all imputed datasets using a score-based logistic regression under an additive model with covariate adjustment for sex, the first three PC eigenvectors for population substructure, and indicator variables for genotyping platform to mitigate potential batch effects. All association analyses were performed using the program SNPTEST^63^. After analysis, variants with regression coefficient of |β|>5 and any erroneous estimates (negative standard errors or *P*-values equal to 0 or 1) were excluded from further analysis.

#### External expression datasets

Expression quantitative trait locus (eQTL) full summary statistics for bulk RNA-seq from 13 human brain regions from the GTEx consortium v8 and v7^64,65^ were downloaded from the GTEx web portal. Donor numbers ranged from 114 (substantia nigra) to 209 (cerebellum). eQTL summary statistics for 8 cortical cell types from single nucleus RNA-seq of 196 donors were downloaded from Zenodo^23^. GWAS summary statistics for Alzheimer disease and Parkinson disease were downloaded from their respective repositories^17,18^. The PSP bulk RNA sequencing data was downloaded from “The Mayo clinic RNAseq study” and the whole blood data was downloaded from the Gene Expression Omnibus portal from a study entitled “Systems-level analysis of peripheral blood gene expression in dementia patients reveals an innate immune response shared across multiple disorders”^26^. All summary statistics were coordinate sorted and indexed with Tabix to allow random access^66^.

#### Fine-mapping

For each locus, we gathered all SNPs within 2-Mb windows (±1Mb flanking the lead GWAS SNP) and filtered out SNPs with a minor allele frequency (MAF)<0.001. We focused on common variants to maximize the relevance of these results to a larger proportion of the PSP population. LD correlation matrices (in units of r) were acquired for each locus from the UK Biobank (UKB) reference panel, pre-calculated by Weissbrod et al.^21^. Any SNPs that could not be identified within the LD reference were necessarily removed from subsequent analyses. Statistical fine-mapping was performed on each locus separately with FINEMAP^20^. Functional fine-mapping was performed using PolyFun + FINEMAP, both of which compute SNP-wise heritability-derived prior probabilities using an L2-regularized extension of stratified-linkage disequilibrium (LD) Score (S-LDSC) regression^21^. For PolyFun + FINEMAP, we used the default UK Biobank baseline model composed of 187 binarized epigenomic and genic annotations^67^. In all subsequent analyses presented here, SNPs that fall within the *MAPT* locus and HLA region/*C4A* locus were excluded due to the particularly complex LD structure^68^. PolyFun + FINEMAP provides a 1) posterior probability (*PP*) that each SNP is causal, on a scale from 0 to 1, and 2) credible sets (CS) of SNPs that have been identified as having a high PP of being causal, which we have set at a threshold of *PP*≥0.95. PolyFun+FINEMAP meets the following criteria: 1) can take into account LD and 2) can operate using only summary statistics. For FINEMAP, we set the maximum number of causal SNPs to five.

#### Cell type-specific epigenomic annotations

For all downstream fine-mapping analyses, we used functional annotations from cell type-specific ChIP-seq annotations of regulatory regions (enhancers and promoters) and cell type-specific DNA interactome anchors from proximity ligation-assisted ChIP-Seq (PLAC-seq)^16^. These same epigenomic datasets are used for both the fine-mapping summary overlap plot and each locus plot and consist of the following cell types: neurons, oligodendrocytes, microglia, and astrocytes. For the fine-mapping summary plot, we also compare the overlap between fine-mapped PSP GWAS SNPs and significant SNPs identified by the HEK293T cell-line MPRA for Alzheimer’s Disease (AD) and PSP^22^. Active promoters and enhancers were defined as follows. H3K4me3 and H3K27ac ChIP-seq data were collected for each purified cell type. Active promoters were defined as the intersection between H3K4me3 peaks and H3K27ac peaks that were within 2 kb of the nearest transcription start site (TSS). Active enhancers were defined as H3K27ac peaks that were not within H3K4me3 peaks.

#### LD-score regression

Stratified LD score regression (S-LDSC) was applied to determine whether specific brain cell type annotations were enriched for heritability of progressive supranuclear palsy^14-16^. Binary annotations were created using active promoters and enhancers, as well as the 1000G Phase 3 panel of common variants that was used in the LDSC baseline annotation model (annotation = 1 if the common variant falls in a promoter/enhancer peak in a particular cell type, annotation = 0 if not)^69^. The cell type-specific enhancer and promoter peak sets were then tested for enrichment of heritability while controlling for the full baseline model.

#### Colocalization and gene prioritization

Two independent pipelines were applied to the GWAS summary statistics to prioritize genes flanking and within the significant loci. We first used the COLOC package to test whether SNPs from different disease GWAS colocalized with expression QTLs from bulk RNA-seq or single-nucleus RNA-seq^70^. For each genome-wide significant locus in the GWAS we extracted the nominal summary statistics of association for all SNPs within 1 Mb either side of the lead SNP (2Mb-wide region total). In each QTL dataset we then extracted all nominal associations for all SNP-gene pairs within that range and tested for colocalization between the GWAS locus and each gene. Where minor allele frequency was missing, we used reference values from the 1000 genomes (phase 3) European superpopulations. Colocalization was performed by comparing the P-value distributions between matching sets of SNPs. To reduce false positives caused by long-range LD contamination, we removed the *MAPT* and *HLA* regions from consideration and restricted locus-gene colocalizations to GWAS-eQTL SNP pairs where the distance between their respective top SNPs was ≤ 500kb or the two lead SNPs were in modest LD (r^2^ > 0.1), taken from the 1000 Genomes (Phase 3) European superpopulations using the LDLinkR package^71^.

Analyses were then performed in GRCh37/hg19 using the INFERNO and SparkINFERNO pipelines which are detailed elsewhere^25,72^. LD-based pruning was run using the 1000 Genome EUR reference genotype panel on all GWS variants (*P*<5×10^-8^, *n*=3016) using r^2^ < 0.7 and 500kb window. This resulted in 108 independent signals (loci). We defined loci to include variants in LD with the tag variant (r^2^≥0.7) restricting to the variants that are at most 1 Mbps away and no more than 1,000 variants between the tag variant and leftmost and rightmost variant in LD with the tag variant. Next, we performed colocalization on each of the 108 loci, against each of the eQTLs from the GTEx v7 dataset^64,70^. In both approaches, we used the posterior probability for colocalization between GWAS and eQTL signals (coloc_PP.H4.abf) at the locus level, as a ranking for causality of each gene at each locus.

#### Imputation of C4A and C4B copy number

*C4* alleles from the genotypes were computed using the HapMap3 CEU reference panel using a protocol generated by Sekar *et al*.^36^ Briefly, VCF files were generated from chromosome six, and imputation was run using BEAGLE^73^. The results were compiled into a table containing *C4A* and *C4B* copy on each subject in the study, except for 16 cases which the program was unable to compute copy number status. Long and short isoforms were not considered in the model. Plink v. 1.90 was run using the same covariates as the main analysis with the addition of the imputed copy numbers for both genes and run separately. Additionally, logistic regression of case control status was run in R comparing *C4A* and *C4B* copy number using the same covariates as the primary association study.

#### Differential gene expression

Raw RNA-seq data from PSP and control postmortem brain was processed using the RAPiD-nf pipeline developed as part of the CommonMind consortium. RAPiD-nf is a pipeline in the NextFlow framework and uses Trimmomatic (version 0.36), STAR (version 2.7a), FASTQC (version 0.11.8), featureCounts (version 1.3.1), and Picard (version 2.20.0) for pre-processing and quality control. RSEM (1.3.1) was used for gene expression estimation^74-77^. After processing, 84 cortical samples and 83 cerebellar samples from the PSP cases were included and 77 cortical and cerebellar samples were used from controls. Principal component analysis on the normalized RNA-seq matrix was performed to identify outliers based on clustering. The RNA-seq matrix was normalized using trimmed mean of M values and transformed using the limma::voom() function and lowly expressed genes removed^78^. Covariates were selected to minimize gene expression differences based on technical variables. Clinical and technical variables from Picard were combined and correlated using variancePartition^79^. Variables that contributed the most to variance in gene expression and had the least overlap with one another were included. Final variables included as covariates were RNA integrity number (RIN), mean insert size, age at death, and biological sex. After normalization and covariate adjustment, differential gene expression (DGE) analysis was performed on 16 genes contained within a 2Mb-wide region flanking each lead SNP using the limma package to compare gene expression of PSP cases and controls^80^. Limma calculated log_2_-fold change, t-statistics, and *P* values for each gene. Because we looked specifically at 16 genes contained within five significant loci, a *P*<0.05/16=0.0025 was considered differentially expressed.

#### Immunohistochemistry

Human brain tissues were fixed in 10% formalin, embedded in paraffin, and cut to a thickness of 6 micrometers (*n*=10 for cases vs. *n*=10 for PSP). Slides were baked and deparaffinized in EZ prep at 72°C for 8 minutes, then pretreated with Heat Induced Epitope Retrieval (HIER) in Tris-EDTA buffer pH 7.8 at 95°C for 64 min in standard cell condition solution one (CC1) using a Ventana Discovery ULTRA (Roche Indianapolis IN). Blocking was then performed in an inhibitor solution for 12 minutes at room temperature. Incubation was then performed with primary antibody oligodendrocyte transcription factor (OLIG2, pre-diluted by the manufacturer) for 40 min at room temperature. A secondary antibody, OmniMap anti-rabbit horseradish peroxidase (HRP), was added for 12 minutes followed by the addition of 3,3′-Diaminobenzidine (DAB) CM / H_2_O_2_ CM with an 8 minute incubation time, and Copper CM was added and incubated for 5 minutes. Next, a denaturation cycle was then run at 95°C for 8 minutes followed by incubation with primary antibody C4a (1:700) for 32 min at room temperature and then with OmniMap secondary anti-Rabbit HRP antibody for 12 minutes followed by purple / H_2_O_2_ incubation at 28 minutes to enhance the bright field color. A denaturation cycle was then run at 95°C for 8 minutes. A final incubation with a third primary antibody against hyperphosphorylated tau (AT8, 1:1500) was run for 32 minutes at room temperature and OmniMap anti-Mouse HRP secondary antibody was added for 12 minutes, followed by GREEN HRP / H_2_O_2_ incubation for 16 minutes and another incubation Green Activator for 16 minutes to enhance visualization. Lastly, a counterstain with hematoxylin was added for 4 minutes, and then a post counterstain Bluing Reagent was added for 4 minutes. A detailed description of the reagents used, and their catalog number can be found in **Supplementary Table 2**.

#### Image Analysis

Five regions containing marked C4a pathology in the white matter on all cases and controls were imaged on a Nikon Eclipse Ci (Nikon Melville, NY) at 20x magnification. The NeuronJ package contained within FIJI v.2.13.1 was used to assess cellular features of complement-activation quantitatively for both the length of the feature and the total number of features^81,82^.

### Biochemical analysis

Western blots were performed using fresh-frozen brain tissues from the prefrontal cortex (*n*=7 for PSP, 6 for controls). Samples were homogenized with a glass-Teflon homogenizer at 500 rpm in 10 volumes (wt/vol) of ice-cold Pierce RIPA buffer (Thermo Fisher Scientific, Waltham, MA) containing Halt protease and phosphatase inhibitor cocktail (Thermo Fisher Scientific, Waltham, MA), incubated on ice for 30 min, centrifuged at 16,000g for 15 min, and then supernatants were collected. For each sample, 30 μg of proteins were boiled in Laemmli sample buffer (Bio-Rad, Hercules, CA) for 5 min, run on 10% PROTEAN TGX Precast Gels (Bio-Rad, Hercules, CA), blotted to nitrocellulose membranes, and stained with C4a antisera (ab170942, 1:1000; Abcam, Waltham, MA). Horseradish peroxidase-labeled secondary anti-rabbit antibody (1:20,000; Vector Labs, Burlingame, CA) was detected by Pierce ECL Western Blotting Substrate (Thermo Fisher Scientific). To quantify and standardize protein levels without reliance on specific housekeeping proteins, total protein was detected with Amido Black (Sigma-Aldrich, St. Louis, MO). Chemiluminescence was measured in a ChemiDoc Imaging System (Bio-Rad, Hercules, CA), and relative optical densities were determined by using AlphaEaseFC software, version 4.0.1 (Alpha Innotech, San Jose, CA), normalized to total protein loaded. *<u>Statistical Analysis</u>*

All non-GWAS were performed in R v4.0 and plotted using ggplot2 v3.4.2. For non-normally distributed data a Wilcox test was used to test for significance, and an ANOVA was used for normally distributed data.

#### Data availability

The publicly available data used here can be found in the following repositories, GTEx web portal (https://gtexportal.org/home/datasets), eQTL single cell data (https://zenodo.org/record/5543735), AD GWAS summary statistics (https://www.niagads.org/datasets/ng00075), PD GWAS summary statistics (https://drive.google.com/drive/folders/10bGj6HfAXgl-JslpI9ZJIL_JIgZyktxn), Mayo clinic RNA seq study (https://adknowledgeportal.synapse.org/Explore/Studies/DetailsPage/StudyDetails?Study=syn5550404), Whole blood microarray data (https://doi.org/10.1101/2019.12.13.875112) the brain plac-seq (https://www.ncbi.nlm.nih.gov/projects/gap/cgi-bin/study.cgi?study_id=phs001373.v2.p2), Picard (https://github.com/broadinstitute/picard/releases), the C4 imputation panel (https://github.com/freeseek/imputec4) and the 1000 genomes reference panel (https://www.internationalgenome.org)

#### Code availability

All software used is publicly available at the URLs or references cited.

## Acknowledgements

This work was supported in part through the computational resources and staff expertise provided by Scientific Computing at the Icahn School of Medicine at Mount Sinai and supported by the Clinical and Translational Science Awards (CTSA) grant UL1TR004419 from the National Center for Advancing Translational Sciences. Research reported in this paper was supported by the Office of Research Infrastructure of the National Institutes of Health under award number S10OD026880 and S10OD030463. The content is solely the responsibility of the authors and does not necessarily represent the official views of the National Institutes of Health.

The authors would like to acknowledge the following tissue repositories for providing the materials necessary to conduct the study: University of Louisville, Australian Brain Bank Network and Flinders University, Barcelona Biobanc and The University of Barcelona, Brain-Net Germany and Neurobiobank Munich, Emory University, Harvard Brain Tissue Resource Center, McLean Brain Bank, Indiana University School of Medicine, Johns Hopkins University, London brain bank, Los Angeles Veterans Association hospital brain bank, Ludwig-Maximilians-Universität München, German Center for Neurodegenerative Diseases (DZNE), Madrid (Universidad Autónoma de Madrid Spain), Massachusetts General Institute for Neurodegenerative Disease, Mayo Clinic Jacksonville, Netherlands Brain Bank and Erasmus University, New York Brain Bank, Columbia University, University of Paris, Southern Texas University, Sun Health Research Institute, University College London Queen Square Institute of NeurologyQueen Square Brain Bank for Neurological Disorders, University of California San Diego, University of California San Francisco Memory and Aging Center, University of Antwerp, University of Michigan, University of Navarra, University of Saskatchewan, University of Southern California, University of Toronto, University of Washington, University of Würzburg, Victorian Brain Bank, Boston University, Emory University, Netherlands Brain Bank and Erasmus University, Oregon Health Sciences University, University of Pittsburgh, University of Miami, University of Washington, University of California Irvine and the NIH Neurobiobank.

## Supplementary Figures and Tables

**Supplementary Table 1.**
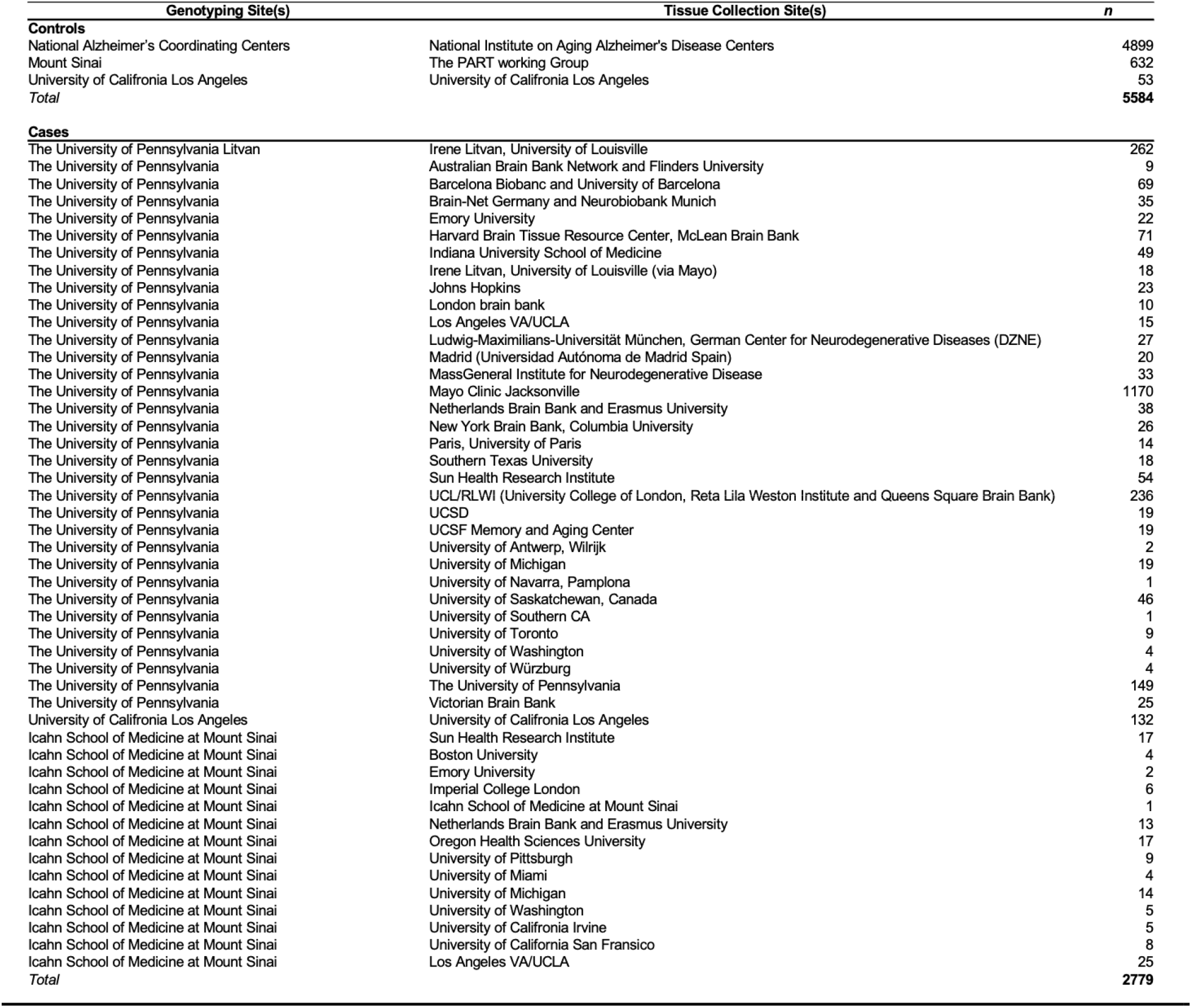
Sources of PSP Cases and Controls.

**Supplementary Table 2.** Detailed description of reagents used in histological staining.

| **PRODUCTS** | **CAT NUMBER** |
| --- | --- |
| DISC. ChromoMap DAB RUO | 760-159 Ventana |
| DISC. Purple Kit RUO | 760-229 Ventana |
| DISCOVERY Green HRP Kit (RUO) | 760-271 Ventana |
| Olig2 Rabbit Monoclonal (EP112) PAb | 760-5050 Cell Marque |
| Recombinant Anti-C4a Rabbit monoclonal [EPR10143] | Ab170942 ABCAM |
| AT8 phospho-Tau Mouse monoclonal(Ser 202, Thr205) | MN1020 Invitrogen |
| DISC. OmniMAP HRP Anti-Rb RUO | 760-4311 Cell Marque |
| DISC. OmniMAP HRP Anti-Ms RUO | 760-4310 Cell Marque |

**Supplementary Figure 1.**
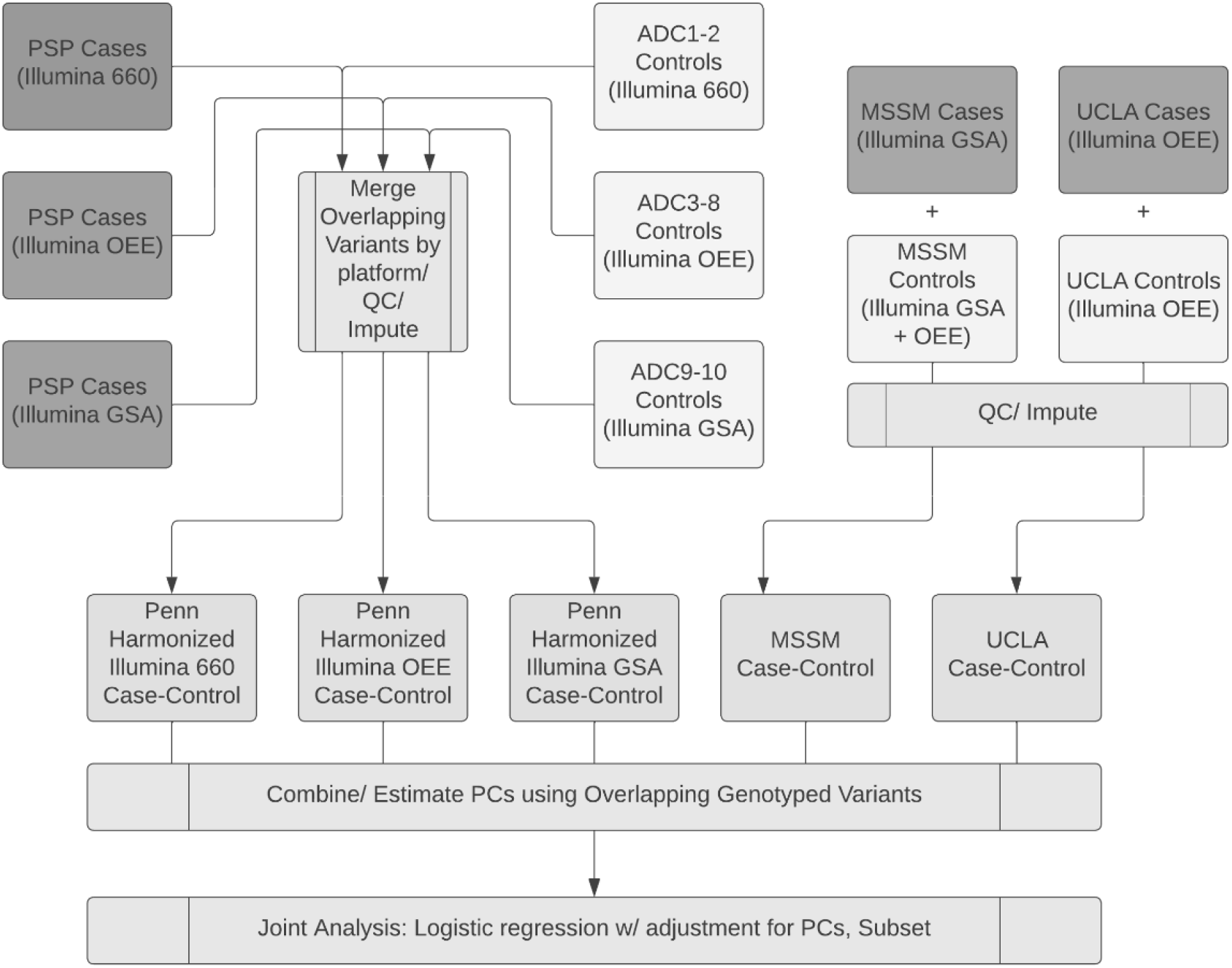
Schematic overview of sample acquisition, genotyping and analysis of progressive supranuclear palsy cases and controls. Global Screening Array (GSA) Omni Express (OEE).

**Supplementary Figure 2.**
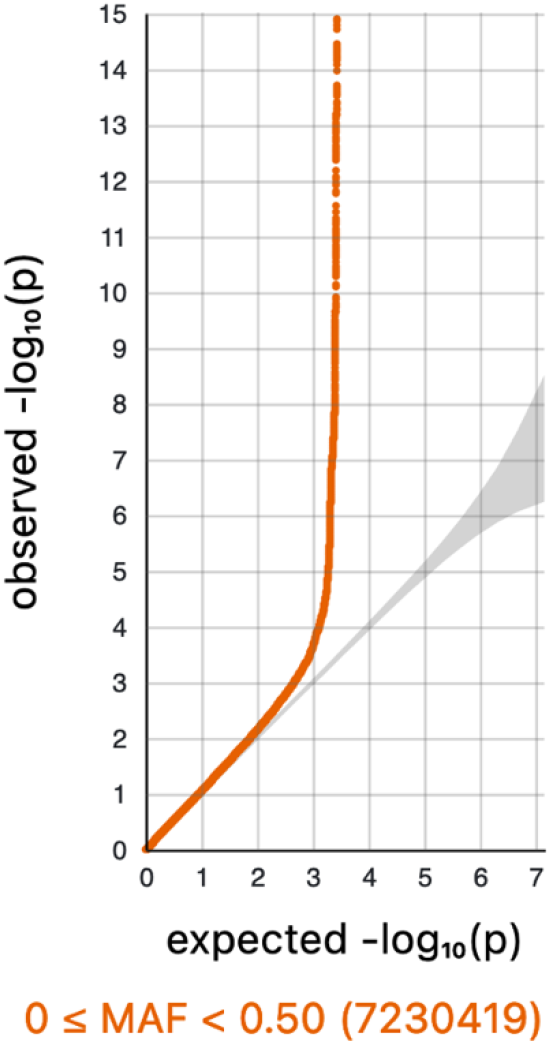
Q-Q Plot of Genome-wide associations with PSP in an analysis of xxx common SNPs.

**Supplementary Figure 3.**
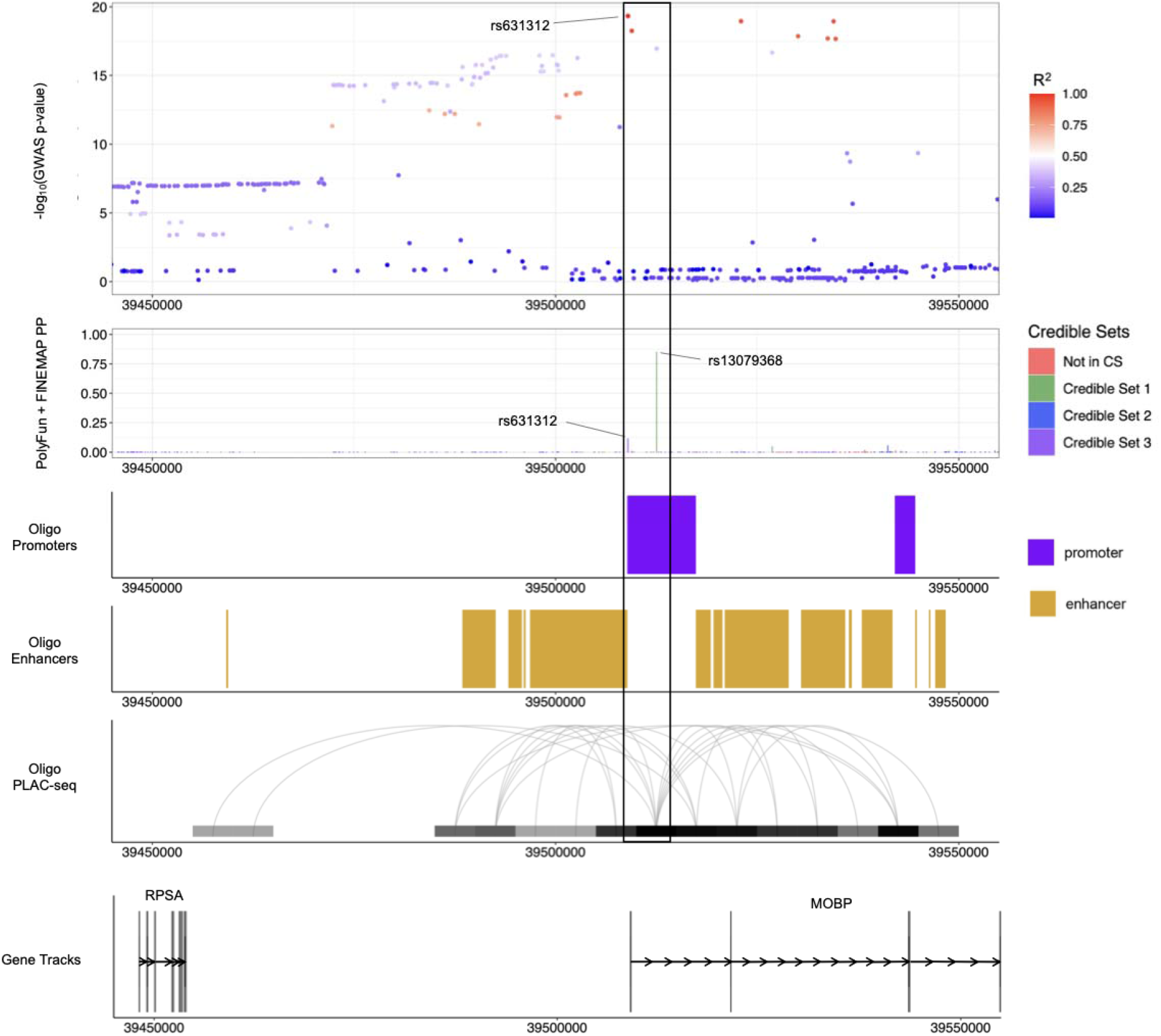
Functional annotations from cell type-specific regulatory regions (enhancers and promoters) and cell type-specific DNA interactome anchors from proximity ligation-assisted ChIP-Seq (PLAC-seq) are shown in the locus containing MOBP SNPs are coloured by their LD with the lead GWAS SNP in Europeans (1000 Genomes European superpopulation).

**Supplementary Figure 4.**
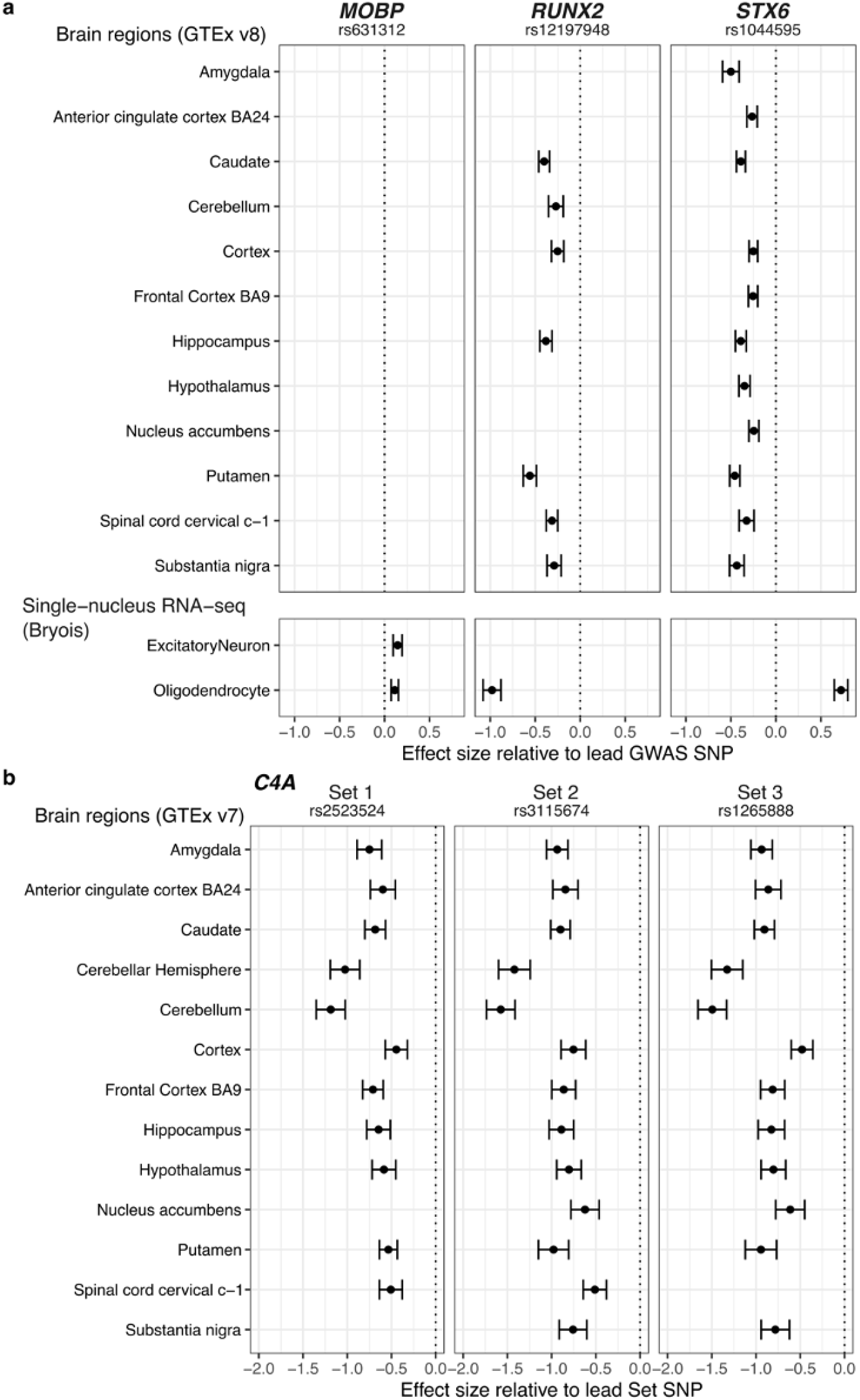
Directions of effect of C4A eQTLs in GTEx brain regions using the lead SNP of the three SNP sets identified by INFERNO in the 6p21.32 locus.

**Supplementary Figure 5.**
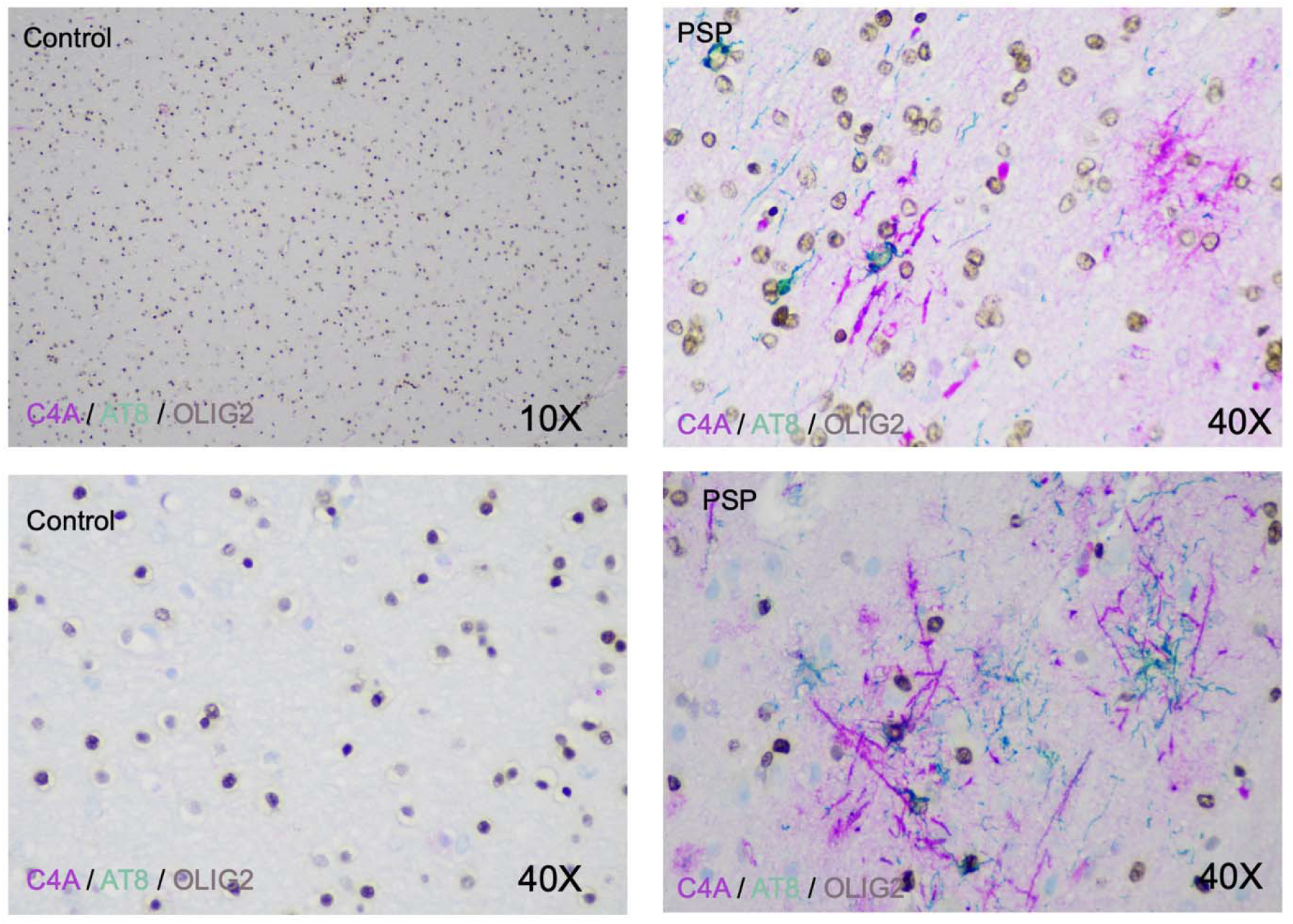
Additional representative staining images of C4A, AT8, and OLIG2 in frontal cortex of human postmortem progressive supranuclear palsy (PSP) and control brain tissue

**Supplementary Figure 6.**
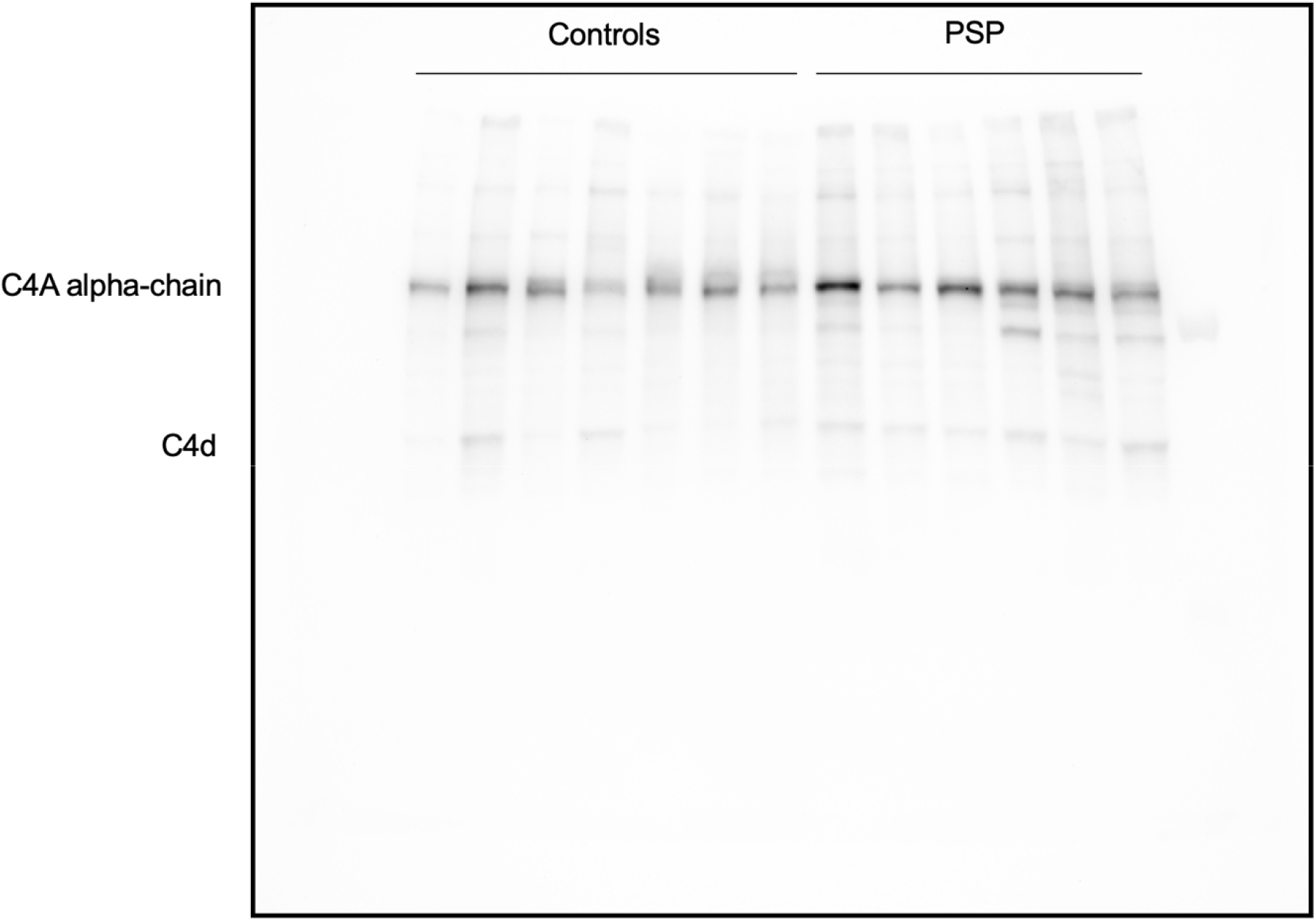
Unedited C4A Western blot of PSP cases and controls

**Supplementary Figure 7.**
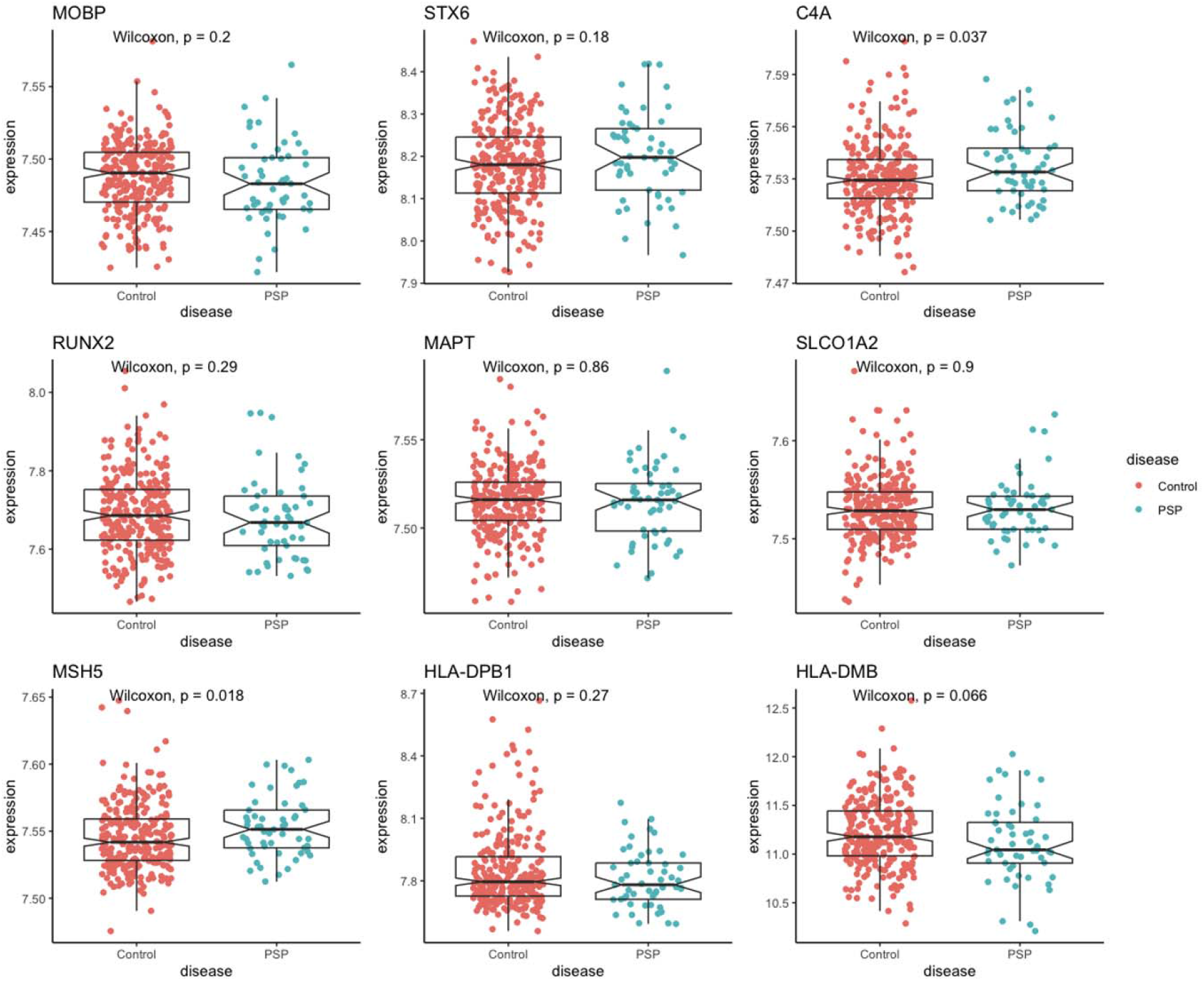
Whole blood gene expression generated from microarray data on genes a select number of genes contained in significant loci.

## Notes

**Funding:** Crary/Farrell Labs: [R01 AG054008, R01 NS095252, R01 AG060961, R01 NS086736, and R01 AG062348 P30 AG066514 to J.F.C. K01 AG070326 and CurePSP 685-2023-06-Pathway to K.F.], the Rainwater Charitable Foundation / Tau Consortium, Karen Strauss Cook Research Scholar Award, Stuart Katz & Dr. Jane Martin. Penn/Lee/Naj/Wang/Schellenberg Labs: [P01 AG017586, U54 NS100693, and UG3 NS104095; RF1 AG074328-01, and P30 AG072979; CurePSP Consortium; Controls were drawn from the ADGC (U01 AG032984, RC2 AG036528), and included samples from the National Cell Repository for Alzheimer’s Disease (NCRAD), which receives government support under a cooperative agreement grant (U24 AG21886) awarded by the National Institute on Aging (NIA). We thank contributors who collected samples used in this study, as well as patients and their families, whose help and participation made this work possible; Control data for this study were prepared, archived, and distributed by the National Institute on Aging Alzheimer’s Disease Data Storage Site (NIAGADS) at the University of Pennsylvania (U24-AG041689); additional salary and analytical support were provided by NIA grants R01 AG054060 and RF1 AG061351] Raj/Humphrey/Ravi: [R56-AG055824, U01-AG068880 U54-NS123743 to J.H., A.R., and T.R.] Goate Lab: [Rainwater Charitable Foundation, NS123746] UCLA/Geschwind lab: [K08AG065519, 3UH3NS104095, Larry L Hillblom Foundation, Tau Consortium] Ross/Dickson: U54 NS100693, P50 AG016574, CurePSP Foundation, Mayo Foundation. Hardy lab: The Dolby Foundation Höglinger Lab: Deutsche Forschungsgemeinschaft (DFG, German Research Foundation) under Germany’s Excellence Strategy within the framework of the Munich Cluster for Systems Neurology (EXC 2145 SyNergy – ID 390857198), DFG (HO2402/18-1 MSAomics), the German Federal Ministry of Education and Research (BMBF, 01KU1403A EpiPD; 01EK1605A HitTau); Niedersächsisches Ministerium für Wissenschaft und Kunst / VolkswagenStiftung (Niedersächsisches Vorab), Petermax-Müller Foundation (Etiology and Therapy of Synucleinopathies and Tauopathies).

**Conflicts of Interest:** AMG is an SAB member for Genentech and Muna Therapeutics. HM consultants for Roche, Aprinoia, AI Therapeutics and Amylyx and is a co-applicant on a patent application PCT/GB2012/052140.

### Competing Interest Statement

AMG is an SAB member for Genentech and Muna Therapeutics.
HM consultants for Roche, Aprinoia, AI Therapeutics and Amylyx and is a co-applicant on a patent application PCT/GB2012/052140.

